# Nascent transcript folding plays a major role in determining RNA polymerase elongation rates

**DOI:** 10.1101/2020.03.05.969709

**Authors:** Tomasz W. Turowski, Elisabeth Petfalski, Benjamin D. Goddard, Sarah L. French, Aleksandra Helwak, David Tollervey

## Abstract

Transcription elongation rates are important for RNA processing, but sequence-specific regulation is poorly understood. We addressed this *in vivo*, analyzing RNAPI in *S.cerevisiae*. Analysis of Miller chromatin spreads and mapping RNAPI using UV crosslinking, revealed a marked 5’ bias and strikingly uneven local polymerase occupancy, indicating substantial variation in transcription speed. Two features of the nascent transcript correlated with RNAPI distribution; folding energy and G+C-content. *In vitro* experiments confirmed that strong RNA structures close to the polymerase promote forward translocation and limit backtracking, whereas high G+C within the transcription bubble slows elongation. We developed a mathematical model for RNAPI elongation, which confirmed the importance of nascent RNA folding in transcription. RNAPI from *S.pombe* was similarly sensitive to transcript folding, as were *S.cerevisiae* RNAPII and RNAPIII. For RNAPII, unstructured RNA, which favors slowed elongation, was associated with faster cotranscriptional splicing and proximal splice site usage indicating regulatory significance for transcript folding.

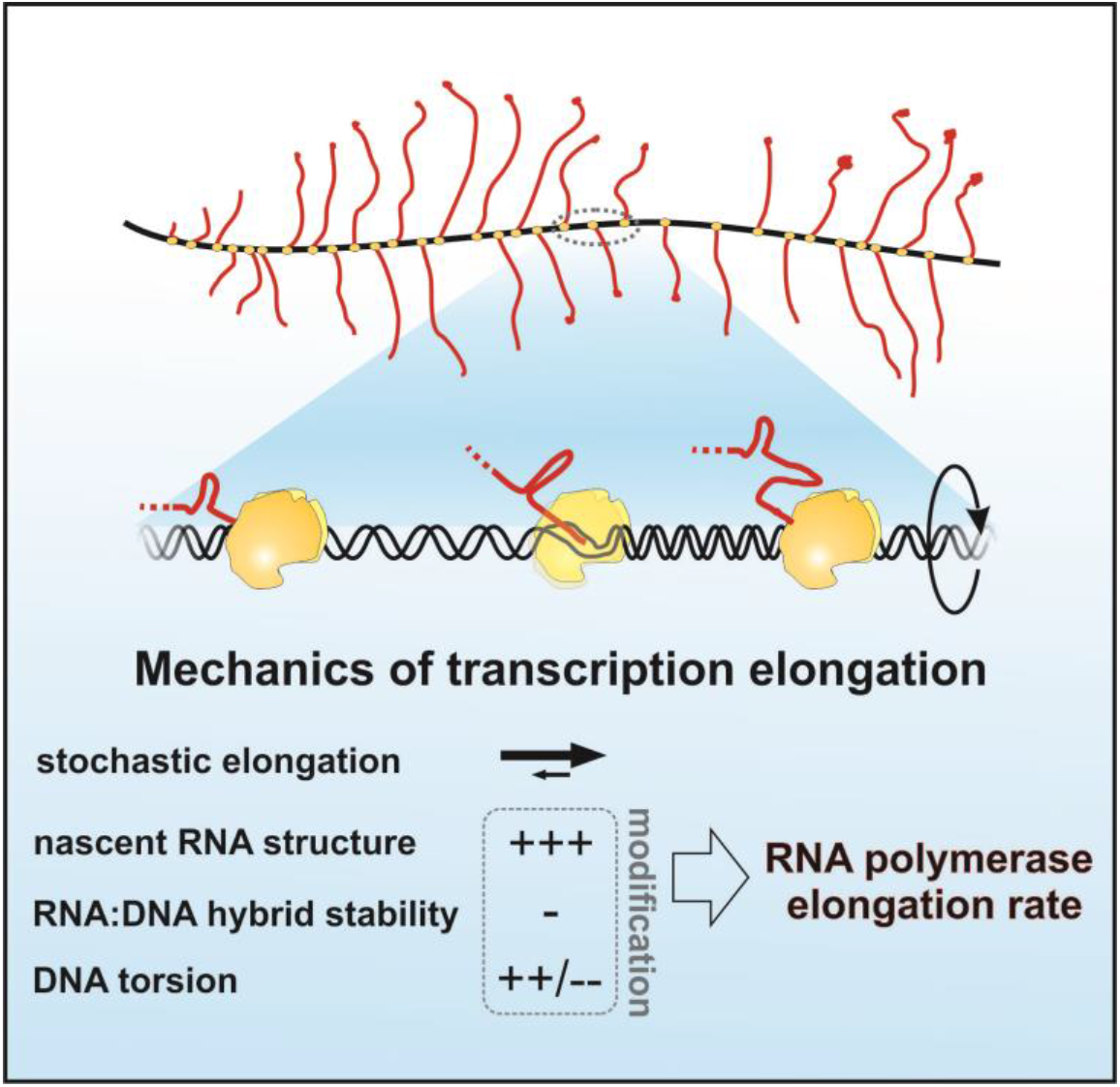

**HIGHLIGHTS:**

Structures in the nascent RNA correlate with rapid elongation by RNAPI *in vivo*
Stable RNA structures limit RNAPI backtracking *in vitro*
GC content in the transcription bubble tunes transcription elongation rate
Nascent transcript folding modulates dynamics of all three RNAPs *in vivo*

## INTRODUCTION

Transcription elongation is composed of many successive cycles of nucleotide addition, in which the translocation step is based on Brownian motion without input of external energy. The major driver of transcription elongation is nucleotide addition, since pyrophosphate release is essentially irreversible, allowing this step to act as a ratchet (Fig. 1A). Dependence on this “Brownian ratchet”, rather than an energy-driven processive mechanism, makes elongation prone to frequent backtracking and potentially sensitive to inhibition or acceleration by quite modest forces (Dangkulwanich et al., 2013). The rate of RNA polymerase (RNAP) elongation can have marked effects on the fate of the newly transcribed RNA; for example, changing RNA folding patterns (Saldi et al., 2018) or the outcome of alternative splicing (Saldi et al., 2016). Deep backtracking is relatively rare compared to the number of nucleotide addition cycles, but in aggregate is widespread in the cell (Sheridan et al., 2019). Notably, RNAPI and III incorporate specific subunits that stimulate endonucleolytic cleavage of nascent RNA when in the backtracked position. In budding yeast these are designated Rpa12 and Rpc11, respectively (Chedin et al., 1998; Kuhn et al., 2007). This is different from RNAPII, in which cleavage of the backtracked transcript is stimulated by an exogenous factor TFIIS (Dst1 in yeast, TCEA1/2/3 in humans). However, the C-terminus of Rpa12 contains a zinc-finger domain homologous to TFIIS (Tafur et al., 2019).

**Figure 1.**
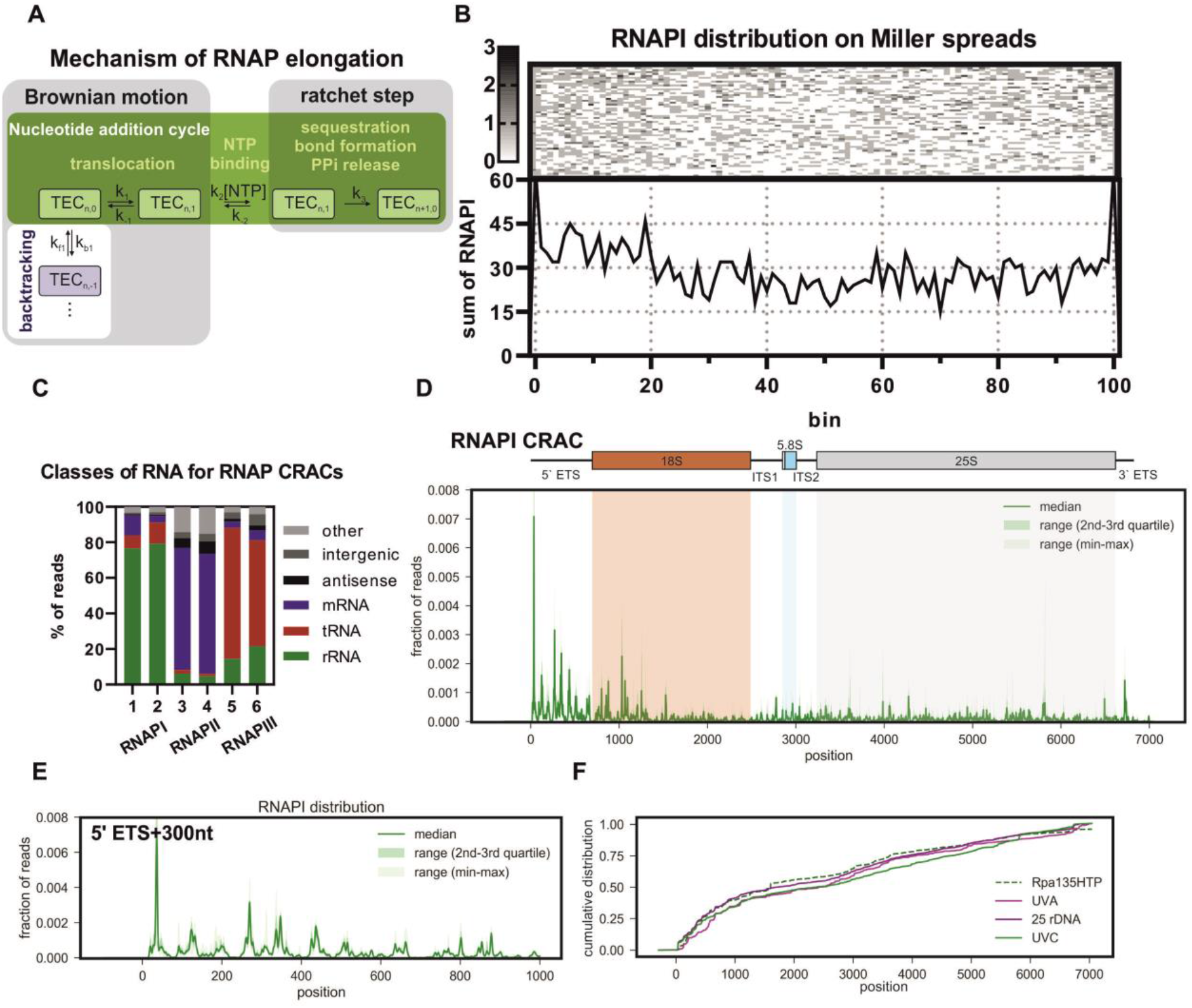
RNAPI distribution along the rDNA *in vivo*. A: Schematic of the Brownian ratchet during the nucleotide addition cycle by the transcription elongation complex (TEC). Translocation of RNAP is driven by Brownian motion, which leads to forward (green background) or backward (white background) movement. RNAP pausing is most frequently associated with the TEC_n-1_ position. Directionality of RNAP is conferred by the ratchet step, where an irreversible phosphodiester bond is formed. Elongation is generated by successive ratchet steps promoted by high NTP concentrations and rapid degradation of the released pyrophosphate (Dangkulwanich et al., 2013). B: RNAPI distribution determined from chromatin spreads. Spreads with 30-70 RNAPI molecules per transcription unit are presented. The first observed RNAPI molecule is assigned to bin 1 and last to bin 100. The plot below is sum of the polymerases in each bin from the genes shown above. C: Transcriptome-wide binding profiles for different RNA classes from replicate CRAC analyses with the catalytic subunits of RNAPI (Rpa190), RNAPII (Rpb1/Rpo21) and RNAPIII (Rpc160/Rpo31). D: Rpa190 CRAC distribution over the *RDN37* gene encoding the pre-rRNA. *Upper panel* Schematic representation of pre-rRNA transcription unit including 18S (red), 5.8S (blue) and 25S (grey) rRNA and <u>e</u>xternal and <u>i</u>nternal <u>t</u>ranscribed <u>s</u>pacers (ETS and ITS respectively). *Lower panel* RNAPI CRAC profile presented as fractions of reads. The solid green line marks the median for six biological replicates, dark green indicates the range between second and third quartiles; light green indicates the range between minimum and maximum values. E: RNAPI CRAC profiles across the first 1000 nt of the transcription unit reveal an uneven distribution, with an apparently regular spacing of peaks. F: Cumulative plot of RNAPI distribution profiles for *RDN37* obtained using CRAC with the second largest subunit (Rpa135-HTP), PAR-CRAC using Rpa190-HTP (UVA), CRAC with Rpa190-HTP in a strain with only 25 rDNA copies (25 rDNA) and in the wild-type (UVC).

The transcription machinery is conserved across all domains of life. In eukaryotes, RNA polymerase RNAPII is most similar to Archaeal RNAP and is therefore believed to be the ancestral form. RNAPI and RNAPIII were derived from RNAPII and specialized for transcription of the pre-rRNA and small non-coding RNAs, respectively (Engel et al., 2018; Khatter et al., 2017; Werner and Grohmann, 2011). Despite functional and structural differences, the basic mechanism of transcription elongation remains the same throughout evolution.

Due to the double stranded, helical structure of DNA, either the DNA or the polymerase must rotate by one complete turn for every 10.5 nt transcribed. In yeast, each active rDNA gene is typically transcribed by ~50 RNAPI molecules, which are associated with nascent pre-ribosomes up to several megadaltons in size. With a transcription rate of ~40 nt sec ^−1^ (Kos and Tollervey, 2010), the transcribing polymerases are predicted to spin the rDNA at ~240 rpm. If all polymerases transcribe at the same rate, there will be no steric strain between adjacent RNAPI molecules. However, any change in the relative positions of transcribing RNAPI molecules generates torsional stress, which will quickly exceed the (low) stalling force of the polymerases (Heberling et al., 2016; Ma et al., 2013; Tantale et al., 2016). The polymerases are therefore torsionally entrained in their relative positions along the rDNA. At the 5’ end, where RNAPI is associate with only a short nascent transcript, we anticipate that torsion can be at least partially released by rotation of the polymerase around the DNA, allowing increased freedom for changes in their relative positions. We therefore predict a gradient of torsional entrainment over the 5’ region of the rDNA. Torsional stress can also be relieved by the action of topoisomerases, Top1 and Top2, which are particularly active on the rDNA reflecting the high transcription rate (Brill et al., 1987; El Hage et al., 2010). However, topoisomerases can unwind a minimum of one complete turn of the DNA, whereas a stalling force is generated by substantially less overwinding for polymerases with spacing typical for the rDNA (120 bp) (Heberling et al., 2016; Ma et al., 2013; Tantale et al., 2016).

The *in vivo* distribution of RNAPI was initially analyzed using Miller chromatin spreads visualized by electron microscopy, in yeast and many other organisms (for an example see (Osheim et al., 2009)). Subsequently, polymerase distributions have been mapped using techniques that include ChIP, NET-seq, CRAC and metabolic labelling (Booth et al., 2016; Churchman and Weissman, 2011; Clarke et al., 2018; Drexler et al., 2019; Mayer et al., 2015; Milligan et al., 2016; Nojima et al., 2015; Schwalb et al., 2016; Turowski et al., 2016; Vinayachandran et al., 2018). Commonly, DNA or RNA is recovered in association with the polymerase and identified by sequencing. The frequency of recovery correlates with the polymerase density at each position. Regions with high signals (peaks) are interpreted as having high polymerase occupancy, and therefore a low elongation rate since RNA transcription is processive process. Conversely, troughs reflect low polymerase occupancy and rapid elongation. Notably, all methods that allow high spatial resolution show markedly uneven polymerase distributions along all genes in yeast and human cells.

Mapping at nucleotide resolution should provide mechanistic information on the process of polymerase elongation. RNAPI is ideally suited to these analyses as it has a high transcription rate, transcribes only the nucleosome-free rDNA and is not known to undergo regulatory phosphorylation (Wittner et al., 2011), facilitating deconvolution of the experimental data. To better understand the mechanism of RNAPI elongation we mapped transcriptionally engaged RNAPI using crosslinking and analysis of cDNAs (CRAC), a method optimized for high specificity of the libraries.

Analysis of the data obtained allowed integration of RNAPI elongation rates with features in the nascent transcript, particularly co-transcriptional RNA folding and GC-content, and with torsional effects generated by DNA unwinding during transcription. We incorporated these results into a kinetic model of RNAPI transcription elongation, providing mechanistic insights into eukaryotic transcription *in vivo*. Due to the evolutionary conservation between polymerases, findings on RNAPI can be applied to other polymerases. Indeed, we demonstrate that our major conclusions on the effects of RNA structure are also applicable to yeast RNAPII and RNAPIII. Moreover, we suggest that modulation of RNAPII elongation by nascent RNA folding is sufficient to affect pre-mRNA splicing, as demonstrated for splicing kinetics and selection of the 3’ splice site in yeast.

## RESULTS

### RNAPI distribution is uneven along the transcription unit

We initially assessed the distribution of RNAPI along the rDNA transcription units using Miller spreads in the wild-type yeast strain (BY4741) growing in YPD medium + 1M sorbitol as previously described (Osheim et al., 2009). To analyze RNAPI distribution we selected 64 spreads for which the full-length rDNA could be unambiguously traced, with polymerases positioned at the 5’ and 3’ ends, and the number of polymerases was around the average number of 50 (range 30 to 70 per rDNA repeat) (see Extended Text for RNAPI quantification). The position of each polymerase along these 64 genes was determined relative to normalized gene length and the results combined into 100 bins (1 bin ≈70 bp, Fig. 1B). Unlike most “textbook” spreads, which generally show even polymerase distribution, but may be highly selected, these data revealed quite heterogeneous distributions, as would be expected for stochastic transcription initiation. The summary plot of RNAPI distribution showed an excess of polymerase density over the 5’ region of the rDNA (Fig. 1B, lower panel). This indicated that the average rate of elongation was lower over the 5’ ETS region, in which major early pre-rRNA assembly events take place (Phipps et al., 2011; Turowski and Tollervey, 2015).

High spatial resolution is not readily obtained using Miller spreads, particularly as the 5’ and 3’ ends can be defined only by the position of the first and last visualized polymerases, and the method is low throughput. To better understand RNAPI distribution, we therefore turned to CRAC, a high resolution, UV crosslinking technique. To perform CRAC, the largest subunit of RNAPI, Rpa190, was genomically tagged with 6xHis-TEV-2xProtA (HTP) and nascent RNA was covalently crosslinked to RNAPI using 254nm UV irradiation, as previously performed for RNAPII and RNAPIII (Milligan et al., 2016; Turowski et al., 2016). After 3-step purification, including stringent, denaturing wash conditions, cDNA libraries were prepared and sequenced using Illumina technology. The CRAC protocol used exclusively recovers RNAs with 3’ hydroxyl groups (see Materials and Methods), expected to represent endogenous 3’ ends of nascent transcripts. Comparison of CRAC data for RNAPI, RNAPII and RNAPIII showed predominant recovery of the relevant subset of RNAs: rRNA for RNAPI, mRNAs for RNAPII and tRNAs for RNAPIII (Fig 1C and Fig S1A).

Qualitative comparison of the CRAC data to Miller spreads revealed a good match in overall profile, confirming the 5’ bias (Figs. 1B, 1D). Average RNAPI density was higher within the first ~1,500 nt, presumably reflecting a lower elongation rate and/or more frequent pausing. This was accompanied by strikingly uneven distribution of read density over this region (Fig. 1E), generating a series of peaks and troughs with apparently regular spacing. Autocorrelation plots (Figs. S1B) confirmed peak separation of around 80 nt, which was very marked over the first 1000 nt.

Highly uneven polymerase distribution was previously observed in datasets for RNAPII and III (Churchman and Weissman, 2011; Milligan et al., 2016; Turowski et al., 2016). However, the 5’ bias in RNAPI distribution and the presence of such distinct peaks were unexpected. We therefore performed extensive validation of the RNAPI CRAC profile using: different crosslinking times; a different RNAPI subunit as bait (Rpa135-HTP); developing photoactivated ribonucleotide (PAR-) CRAC, based on UVA irradiation and 4-thiouracil labeling; and strains with a decreased number of rDNA repeats (25 rDNA) (Fig. S1; see detailed description in Extended Text). Notably, all of these analyses yielded RNAPI distributions that were consistent with the results of CRAC with Rpa190 (Fig. 1F). Further analysis was based on the median of six biological replicates, using Rpa190-HTP and UVC (254nm) crosslinking (Fig. S1K).

The strong, 5’ peak of RNAPI density was centered around +36 (Fig. 1E). The reported RNAPI footprint is ~38 nt, so this is the position expected for a polymerase immediately adjacent to another RNAPI, initiating at +1. We therefore speculate that the +36 peak reflects RNAPI that remains in an initiation state (Engel et al., 2017). Release into an elongation state is expected to increase the elongation rate and might be associated with re-arrangements within the polymerase. In subsequent analyses, we will not consider the 5’ peak, but will focus on elongation steps during RNAPI transcription. We note, however, that this prominent peak would increase the accuracy of 5’ end positioning in the Miller spreads.

### RNAPI density correlates with features in the nascent pre-rRNA

Conceivably, spacing of the 5’ ETS peaks might correspond to the size of the polymerase itself. However, the footprint of RNAPI was determined by cryo-EM to be 38 nt, while the minimal distance between the polymerases on the transcription unit is only slightly longer, as determined by cryo-EM tomography (Engel et al., 2013; Neyer et al., 2016; Tafur et al., 2016). This made it unlikely that the ~80 nt peak separation corresponded to “pileups” of polymerases in close proximity. This would also not be consistent with the EM data, in which any pileups would be visualized (El Hage et al., 2010).

We also considered that the distribution of RNAPI might be influenced by chromatin structure, as found for RNAPII (Churchman and Weissman, 2011; Milligan et al., 2016). The actively transcribed rDNA repeats are not packaged into nucleosomes, but are associated with the DNA binding protein Hmo1, which is related to human HMG1 (Hall et al., 2006; Merz et al., 2008; Wittner et al., 2011). Rpa190 CRAC performed in an *hmo1*Δ strain still showed a 5’ bias and stable peaks over the 5’ region of the rDNA, indicating that these do not reflect DNA packaging by Hmo1 (Fig. S2A-B).

#### High GC content moderates the elongation rate of RNAPI

We next assessed whether features in the nascent pre-rRNA could affect RNAPI elongation kinetics. A short RNA:DNA hybrid is present in the transcription bubble within the RNA polymerase elongation complex (Fig. 2A). For human RNAPII, stable RNA:DNA hybrids in the transcription bubble are more frequently associated with paused/backtracked states (Lukačišin et al., 2017; Schwalb et al., 2016). We used a peak-calling algorithm to define peaks and troughs in the RNAPI density (for an example region see Fig. S2C), and then determined G+C content around each feature (peak or trough). Since the reads are 3’ mapped, the read density indicates the positions of 3’ ends of nascent transcripts within RNAPI. The 10 nt sequence immediately upstream from the feature corresponds to the RNA:DNA hybrid forming the transcription bubble (see Fig. S2D for a schematic). This region showed a higher percentage of G+C for peaks than for troughs (transc. bubble in Fig. 2B). This correlation was seen considering either the entire rDNA (*RDN37*, p<8·10^−5^) or the 5’ ETS alone (p<5·10^−3^). As a control, we determined the G+C content for the combined region 10 nt upstream plus 10 nt downstream from each peak and trough (control in Fig. 2B, p>>0.05), which showed no significant differences. An additional possibility was that unwinding of the template DNA in front of the transcription bubble could be slowed by a high G+C content. However, analysis of the first 10 nt downstream from features showed no clear correlation between peaks of RNAPI density and elevated G+C content, or even an opposing trend (Fig. S2E, p=5·10^−4^ for *RDN37* and p>>0.05 for 5’ ETS). Overall, the data support the model that elevated G+C content in the RNA:DNA hybrid within the transcription bubble is associated with increased RNAPI occupancy, presumably reflecting slowed or transiently paused RNAPI.

**Figure 2.**
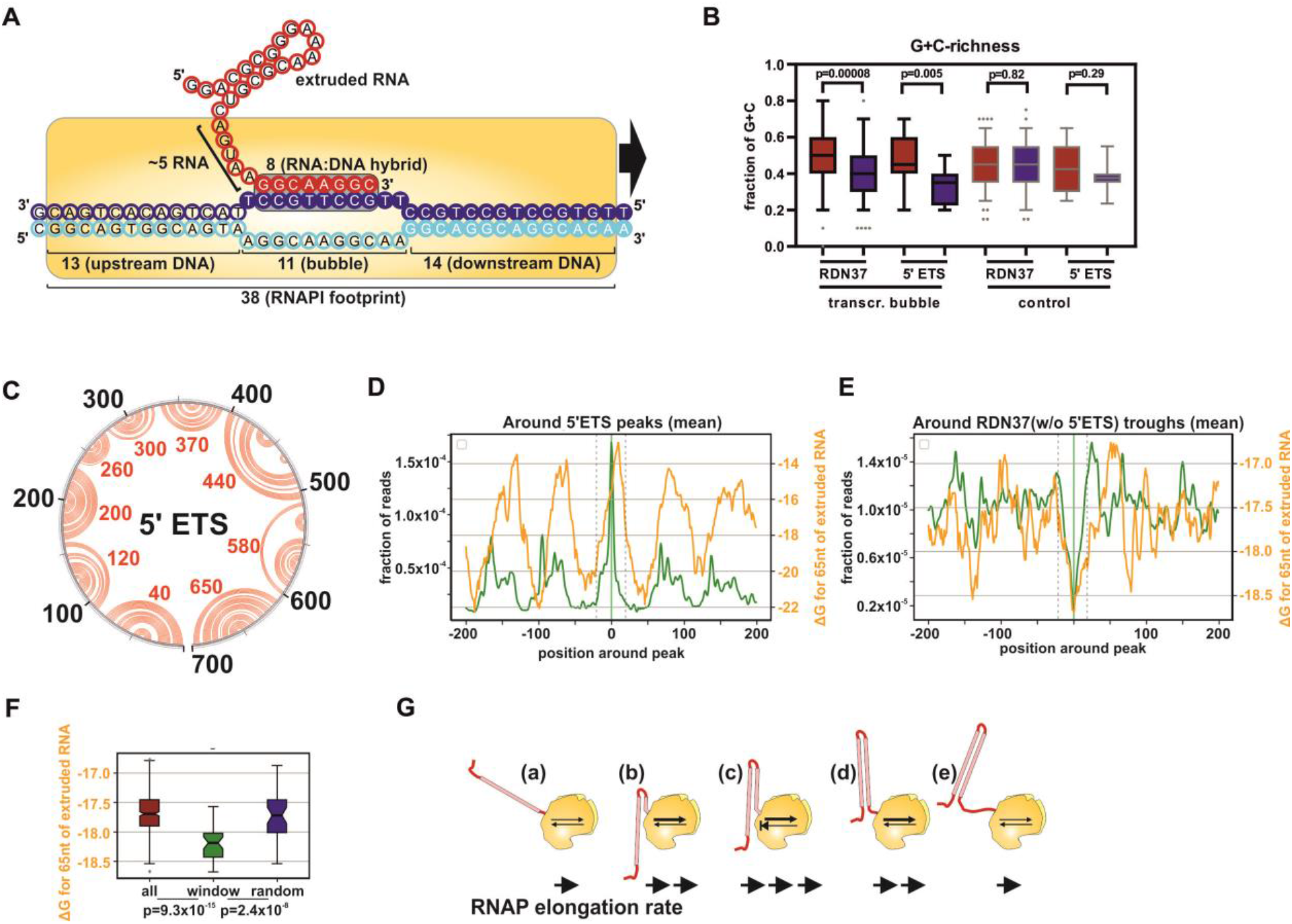
RNAPI density correlates with features in the nascent pre-rRNA. A: Schematic representation of RNAPI transcription, including lengths of DNA and RNA elements hidden in the complex. A 8bp RNA:DNA hybrid forms the transcription bubble, 13nt of RNA is buried in the RNAPI complex and 38bp of DNA is covered by RNAPI. Data derived from the structure of the RNAPI elongation complex PDB: 5M5X (Tafur et al., 2016). B: G+C-content of RNA:DNA hybrids. Boxplot presenting the distribution of G+C-content among peaks (red) and troughs (blue) within *RDN37* or the 5’ ETS. G+C content is calculated for 10 nt upstream from each feature (transcr. bubble) or 10 nt upstream plus 10 nt downstream from each feature (control). C: Secondary structure of the 5’ ETS. *In silico* prediction of UNAfold package (http://unafold.rna.albany.edu/). The black circle represents positions within 5’ ETS, red arches show predicted interactions between bases. Red numbers indicate the loop position for each hairpin. The structure of the 700 nt long, 5’ ETS shows strong, regular stems. D: RNAPI CRAC peak metaplot for the 5’ ETS with the folding energy of nascent transcript. Folding energy (ΔG in Kcal mol^−1^) of 65 nt behind RNAPI was predicted and offset by 15 nt. Stronger structures have lower ΔG. E: RNAPI CRAC trough metaplot for *RDN37* without the 5’ ETS, with folding energy of the nascent transcript. Folding energy as (D). The area between dashed grey lines was used as the window for boxplot (F). F: Boxplots for (E) comparing distribution of folding energy. Full 200 nt window (all), 40 nt region within between dashed grey lines in panel E (window) or random 40 nt (random). P-values were calculated using a two-sided T-test. G: Model: Structures forming in the nascent RNA promote transcription elongation by limiting translocation backward and therefore increasing the elongation rate.

#### Folding of the nascent RNA promotes RNAPI elongation

The yeast 5’ ETS folds into ten stable, extended hairpin structures (Sun et al., 2017) (Fig. 2C). To examine the influence of the RNA structure forming just behind RNAPI, we initially calculated the folding energy for a rolling window of 80 nt upstream from each nucleotide position in the pre-rRNA, which is similar to the average length of 5’ ETS hairpins. Comparison with the RNAPI CRAC peaks showed an apparent correlation with the predicted folding energy across the 5’ ETS (Fig. S2F, R^spearman^=0.65; as a window of 80 nt is used, the folding energy line commences at +80).

To more systematically compare folding to RNAPI density, we used peak and trough metaplots (Fig. S1F). In Fig. S2, the zero position represents the maximum (Figs. S2G and S2H) or minimum (Fig. S2I) for the sum of all peaks or troughs identified by the peak-calling algorithm. This revealed a striking correlation, in which peaks of RNAPI density were associated with weak structures in the nascent pre-rRNA, especially over the 5’ ETS (Figs. S2G, R^spearman^=0.78 and 2D R^spearman^=0.53; structures are plotted as ΔG, with lower values representing greater stability). Conversely, regions of low RNAPI occupancy were correlated with stable structures in the nascent transcript (Fig. S2I). Each position on the x axis shows the average folding energy for the nascent transcripts associated with all polymerases located at that distance from the peak (or trough).

To better understand the relationship between pre-rRNA folding and elongation, the analysis was repeated using a range of window sizes to calculate folding energy. In addition, an “offset” was added, since the terminal ~15 nt of the transcript is located within the polymerase and unable to participate in folding (Figs. S2J). The data indicated that the best correlation was generated by using 65 nt of RNA to calculate folding with a 15 nt offset. The correlation was most marked over the 5’ ETS region (Fig. 2D) but was also observed when the *RDN37* gene was analyzed excluding the 5’ ETS (Figs. 2E and 2F, p<10^−7^).

We conclude that weak structures in the nascent pre-rRNA behind the RNAPI coincide with sites of slowed elongation (high RNAPI density), whereas strong pre-RNA structures correlate with rapid elongation (low RNAPI density).

Since elongation is driven by Brownian motion, there is the potential for backtracking prior to each nucleotide addition step (see Fig. 1A). During backtracking, the newly synthesized region of the nascent transcript must re-enter the exit channel of the polymerase. Only single stranded RNA can enter the polymerase, indicating that backtracking should be strongly opposed by formation of stable RNA structure in the nascent transcript. Moreover, there is a decrease in free energy, i.e. an increase in the structure stability as each additional base pair is formed in extended stems, which might also favor elongation over backtracking. We therefore postulate that stable cotranscriptional folding of the nascent pre-rRNA strongly promotes transcription elongation *in vivo* (Fig. 2G). A consistent observation has been made using single-molecule, *in vitro* transcription assays with yeast RNAPII and mitochondrial RNAP (Zamft et al., 2012).

We note that the 5’ ETS has very stable overall folding (ΔG -265 Kcal mol^−1^ over 700 nt) relative to the 5’ region of the 18S rRNA (ΔG --220 Kcal mol^−1^ over first 700 nt) despite having a low G+C content (see Discussion). This suggests that structure in the 5’ ETS has been selected, possibly to promote elongation.

### RNA structures limit RNAPI backtracking *in vitro*

Determination of nascent structures *in vivo* remains beyond current methodology, since structure probing requires chemical treatment for minutes, whereas the effects of nascent RNA are expected over 1-2 sec due to the fast elongation rate of RNAPI (~40nt sec^−1^). To validate the conclusion that the structure in the nascent pre-rRNA limits backtracking, we therefore used an *in vitro* RNAPI transcription system (Pilsl et al., 2016). In this, immobilized RNAPI binds an RNA:DNA scaffold, which mimics the transcription bubble, and elongates the transcript following nucleotide addition. The products are gel separated and visualized using a fluorescent label on the RNA primer (Fig. 3A). Within RNAPI, Rpa12 specifically stimulates endonuclease cleavage of nascent RNA in the backtracked position (Kuhn et al., 2007). Backtracking therefore leads to truncation of previously elongated pre-rRNA transcripts.

**Figure 3.**
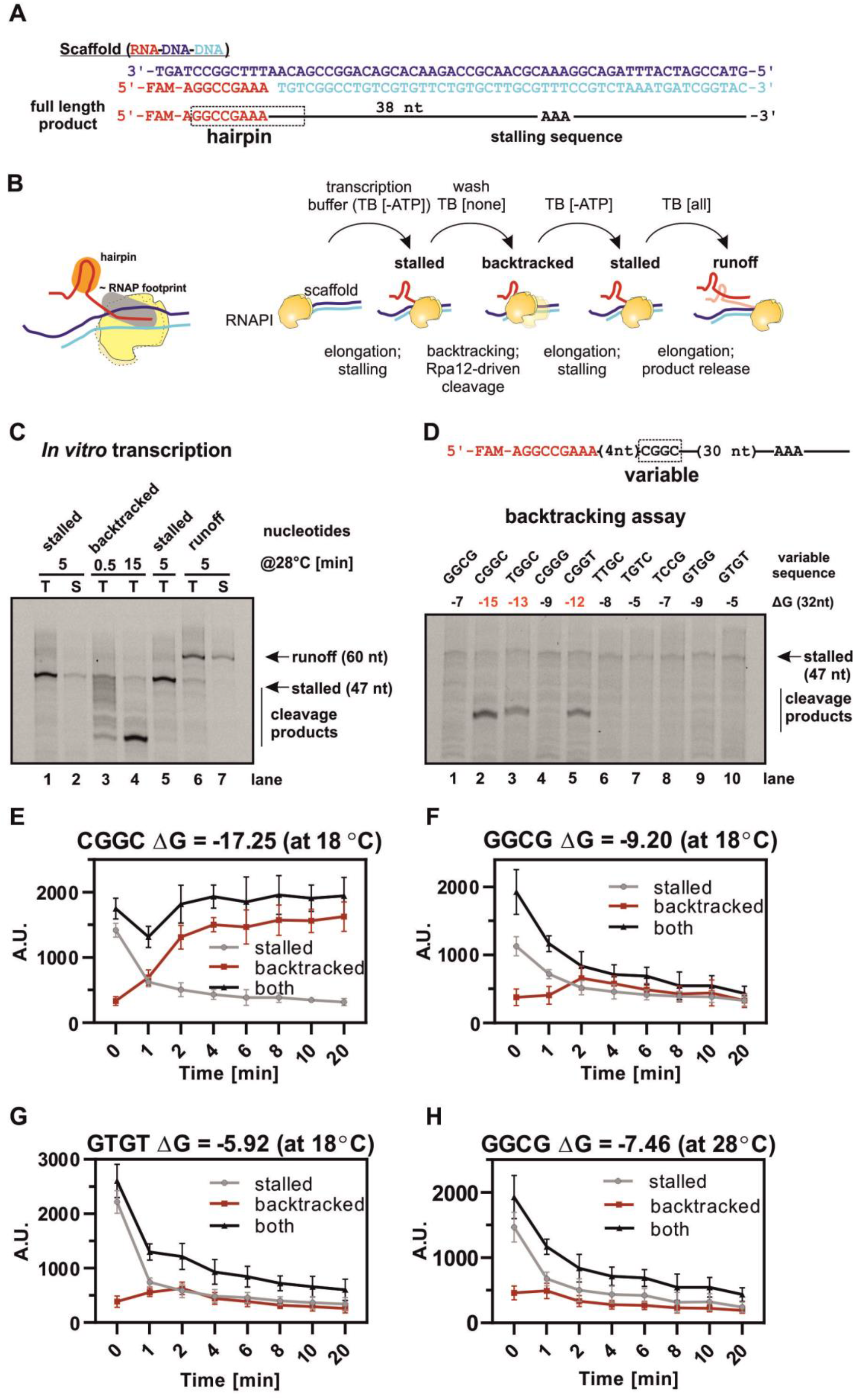
Strong structures in the nascent RNA limit RNAPI backtracking and promote elongation *in vitro*. A: Sequence of RNA-DNA-DNA scaffold and schematic of the 60 nt RNA product with the stalling sequence marked (AAA). B: Schematic of the *in vitro* assay. RNAPI elongation complex (yellow), DNA (light and dark blue) and RNA (red) with the hairpin structure highlighted (orange). [-ATP] - transcription buffer (TB) without ATP to induce stalling (“stalled”); [none] – incubation in TB without nucleotides to promote backtracking and Rpa12-driven cleavage (“backtracked”); [all] - TB supplemented with all NTPs to generate runoff products (“runoff”). C: *In vitro* assay performed according to scheme in (B). T - total, S - supernatant only. The experiment was performed in one tube and aliquots were taken for each condition. D: *In vitro* assay of RNAPI backtracking. *Upper panel:* Schematic of the 60 nt RNA product with four variable nucleotides and stalling sequence (AAA). *Lower panel:* Set of ten scaffolds with different folding energies of nascent RNA, transcribed in TB without ATP to induce stalling, washed and incubated for 15 min at 28°C to allow for Rpa12-driven cleavage. Only sequences with the highest stability limited RNAPI backtracking and produced prominent bands. E – H: Structured RNA reduces back-translocation *in vitro*. Assay performed as panel D. Quantification of stalled peaks, backtracked peaks and the sum of both. Variable scaffold sequence, incubation temperature and predicted folding energy are marked above each plot.

RNAPI was purified via Rpa135-HTP and bound to IgG-conjugated magnetic beads to allow rapid exchange of transcription buffer. Nascent transcripts are retained on the beads in association with the polymerase. The template DNA included a sequence that generates a stem-loop structure in the RNA, close to the 5’ end of the transcript. The transcript lacked A residues other than a sequence of three adenines (AAA) close to the 3’ end of the template (Figs. 3A and 3B). Incubation for 5 min at 28°C in the presence of nucleotides (GTP, UTP, CTP) without ATP [-ATP] resulted in transcription elongation and stalling at the AAA sequence (“stalled”) (Figs. 3B and 3C, lanes 1 and 2). Nucleotides were washed out and the elongation complex was incubated for 15 min at 28°C to allow for RNAPI backtracking (“backtracked”). This generated shorter products, observed as a smear on the gel (Fig. 3C, lane 3-4). These are due to Rpa12 cleavage of the backtracked transcript, as shown by their absence when the same assay was performed using RNAPI purified from a Rpa12ΔC strain (Lisica et al., 2016), in which Rpa12 lacked the C-terminal domain required for cleavage (Fig. S3A, lanes 8 and 9).

Cleavage by Rpa12 should reposition the 3’ end of the nascent transcript in the active site (Lisica et al., 2016; Prescott et al., 2004). Consistent with this expectation, we were able to restart transcription elongation by nucleotide re-addition. Addition of buffer lacking only ATP [-ATP] regenerates the stall (“stalled”), whereas addition of all four nucleotides [all] generates the full-length transcript (“runoff”) (Fig. 3C, lane 5-7). The full-length runoff product was released by RNAPI into the supernatant fraction (Fig. 3C, lane 7).

To compare sequences with different folding energy, we designed *in silico* a construct with four random nucleotides (Fig. 3D, *upper panel*). The predicted folding energy of the stalled, nascent transcript was calculated, and we selected ten sequences for experimental analysis, with a range of stabilities (ΔG –5 to –15 Kcal mol^−1^ at 28°C; low ΔG corresponds to greater stability). In the backtracking assay, samples were first incubated in [-ATP] transcription buffer to induce stalling, then washed and incubated without nucleotides [none] for 15 min at 28°C to allow for RNAPI backtracking (Fig. 3D, *lower panel*). Among the ten constructs tested, only three generated clear, stabilized cleavage products (Fig. 3D, lanes 2, 3 and 5). Notably, these correspond to nascent RNAs with the most stable structures (ΔG –12 to –15 Kcal mol^−1^). We predict that this represents the strength of RNA structure needed to efficiently block further backtracking. Moreover, the cleavage product was more abundant for the construct with ΔG – 15, than for the constructs with ΔG –12 or –13. These results confirm that stable structures in nascent RNA limit backtracking by RNAPI.

Weaker structures did not generate stable stalls at 28°C, but might still affect RNAPI back translocation. To assess this, the RNAPI backtracking assay was analyzed at short time points, and with reduced temperature to slow the polymerase (Fig. 3E-H).

The folding energies of scaffolds used in Fig. 3D were recalculated for 18°C. For the strongest hairpin CGGC (ΔG -15 at 28°C; ΔG -17 at 18°C) we observed very rapid backtracking even at 18°C (Figs. 3E; S3B). By 2 min nearly all RNAPI complexes were lost from the stalled position and accumulated in backtracked positions stabilized by the 5’ terminal stem. These complexes were then stable for at least 20 min of incubation.

We next tested two hairpins that did not generate stable products at 28°C; GGCG (ΔG -7 at 28°C: ΔG -9 at 18°C) and GUGU (ΔG -5 at 28°C; ΔG -6 at 18°C). At 18°C, both transcripts generated a clear but transient gel band corresponding to backtracked RNAPI, which was most prominent at 2 min and destabilized during longer incubation (Figs. 3FG, S3DE). This was more persistent for the more stable GGCG transcript than for GUGU. We also tested the GGCG transcript over a time course at 28°C (Fig. 3H and S3F). The backtracked peak was reduced at 28°C but was still observed after 10 min of incubation, and produced an RNA shortened to 6 nt (Fig. S3G).

Altogether, these kinetic assays revealed that strong structures block backtracking, whereas weaker structures slow the kinetics of back translocation, proportionally to their stability.

### Mathematical model of RNAPI transcription

The CRAC experiments are an average of many individual UV crosslinking events, each reflecting the state of one RNAPI molecule out of the billions present in the culture. For each RNAPI multiple factors influence its behavior. To better understand the contributions of the different components to overall transcription, we developed a mathematical model for RNAPI transcription. Briefly, the model is based on simulations of individual RNAPI molecules initiating and transcribing a 7,000 nt RNA. The key parameters of the model include the (stochastic) initiation frequency and the probability of forward or reverse translocation. The latter is influenced by a several factors: 1) The effects of DNA torsion on the probability of elongation versus backtracking; 2) The effects of structure in the nascent transcript; and 3) the stability of RNA-DNA the duplex in the transcription bubble.

The parameters are briefly described below and discussed in more detail in the Extended Text (Figs. 4 and S4). In this section, “RNAP” is used for statements universal to all RNA polymerases and “RNAPI” for features specific to RNA polymerase I.

**Figure 4.**
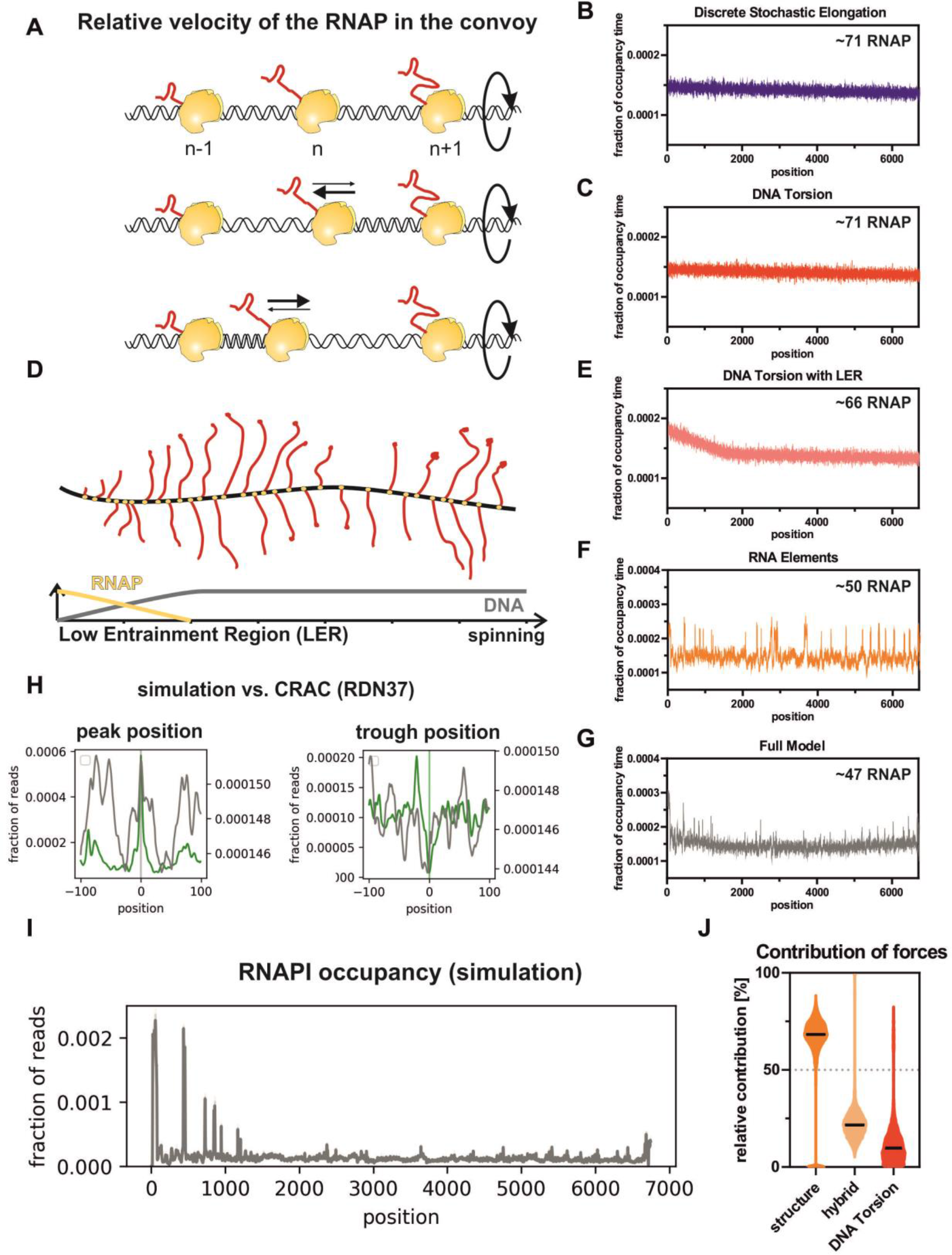
Mathematical model of RNAPI transcription. A: Schematic of polymerase convoys, in which a group of RNAP complexes moves along the rDNA transcription unit while DNA screws through the polymerases. In this model the distance between initially loaded RNAP molecules is retained by the DNA helix, which behaves as an elastic rod and generates force when over- or under-wound. B: Modeled RNAPI occupancy along the transcription unit using model of stochastic initiation and discrete, stochastic elongation. The average number of RNAP molecules per transcription unit is indicated. C: Modeled RNAPI occupancy for the DNA Torsion model. D: Schematic representation of Low Entrainment Region (LER). RNAPI molecules are initially able to rotate around the DNA, allowing changes in their relative positions without generating torsion. They become progressively entrained by torsion, which is fully engaged at around +2Kb. Black line; degree of torsional entrainment. Yellow line; ability of RNAPI to reduce torsion by rotation around the rDNA. E: Modeled RNAPI occupancy for the DNA Torsion model including a LER of 2000 nt. DNA Torsion is engaged linearly between positions 0 and 2000. F: RNAPI occupancy for the model with RNA Elements. Translocation is modified positively by structure in the nascent RNA extruded from the polymerase and negatively by the stability of RNA:DNA hybrid within the transcription bubble. G: Full Model, including DNA Torsion, Low Entrainment Region and RNA elements. This recapitulates experimental data to the greatest extent. Areas that do not correlate with experimental data are predicted to be affected by trans-acting factors binding co-transcriptionally (see Discussion). In panels B-F: The average of RNAPI occupancy for 64 simulations is presented. 200 time points were collected for each 2,000 sec simulation. H: RNAPI CRAC peak and trough metaplots together with simulated data. Full Model (in grey) relative to CRAC data (in green). I: Modeled RNAPI occupancy plot generated using the Full Model. Four datasets of 256 simulations were used to generate profiles. *In silico* “reads” were sampled from interpolated profiles and processed in the same way as RNAPI CRAC data. J: Violin plot of factor contributions at each elongation step within the full model.

#### Starting premises

##### 1. Stochastic initiation events

Yeast cells contain ~200,000 ribosomes, which must be doubled each generation (100 min); corresponding to ~2,000 pre-rRNAs transcribed to completion per minute. There are ~150 rDNA repeats, of which ~50% are transcriptionally active (Dammann et al., 1993). From these numbers we predict that at least one 7 kb-long, 35S pre--rRNA should be fully transcribed from each active rDNA every three seconds. We therefore tested rates of stochastic initiation over a range from 0.33 to 1.0 sec^−1^, limited by the requirement that the preceding RNAPI has cleared the initiation region. A mean stochastic initiation rate of 0.8 sec^−1^ generated RNAPI loading similar to that observed with Miller spreads (~50 per rDNA unit) (Fig. S4F).

##### 2. Stochastic elongation

The reported, average *in vivo* transcription rate, across the entire yeast 35S pre-rRNA, is ~40nt sec^−1^, based on kinetic*, in vivo* labeling and modeling (Kos and Tollervey, 2010) and our calculations support this figure (for details see Extended Text). However, this apparent velocity is generated by the sum of multiple stochastic events. As indicated in Fig. 1A, elongation is dependent on Brownian motion, which can result in forward (elongation) or reverse (backtracking) movement. Analysis of elongation *in vitro* identified the distribution of nucleotide incorporation (elongation) rates at a nucleotide level (Adelman et al., 2002). These rates directly reflect the range of time delays before elongation or backtracking. Note that although transcription elongation rates are often described as a velocity, the polymerase does not have momentum and the time delay for each translocation event is independent and stochastic. At each time step in the model, the probability of translocation is random, chosen from a distribution derived from the experimental data. The sum of these discrete stochastic delays creates the measured transcription rate. Together with stochastic initiation, this generated a model for the distribution of RNAP termed “Stochastic Elongation”.

##### 3. Effects of DNA torsion

During transcription the DNA or the RNAP plus nascent transcript must rotate through 360° for each 10.5 nt incorporated (see Fig. 4A). The mass of all polymerases plus nascent pre-ribosomes is very much greater than that of the rDNA, and it seems very likely that the latter is rotated during pre-rRNA synthesis in agreement with the model of immobilized RNAP (Iborra et al., 1996). If all RNAP molecules move in synchrony the torque from each will be equal, so no torsional stress will accumulate between adjacent polymerases. However, alterations in relative positions will result in positive supercoils between approaching polymerases, and negative supercoils between separating polymerases (Fig. 4A). Positive supercoils in front of the polymerase overwind DNA and strongly resist strand opening during elongation. Even modest changes in relative position can exert sufficient force to effectively block transcription elongation *in vitro* (Heberling et al., 2016; Kim et al., 2019; Lesne et al., 2018; Tantale et al., 2016). Conversely, negative supercoiling in front of RNAP causes underwinding and promotes translocation. Similarly, positive supercoils behind the polymerase resist backtracking, while negative supercoils promote it. The torque generated by torsion acts as an elastic rod, resulting in torsional entrainment of relative RNAP separation.

To reflect torsional entrainment, changes in the relative positions of adjacent polymerases invoke an alteration in the time distribution for their elongation steps (Fig. 4A); An *n* polymerase that approaches the flanking n+1 polymerase is less likely to rapidly translocate (positive supercoils in front), whereas an *n* polymerase that approaches the n-1 polymerase is more likely to translocate (positive supercoils behind). The effect on elongation of this torque assisted motion is included in the model as “DNA Torsion”.

Topoisomerases can relax positive or negative supercoils and are necessary to maintain transcription of the rDNA (Brill et al., 1987; El Hage et al., 2010). The full model included the potential for topoisomerase activity to release accumulated torsion, dependent on RNAPI separation distance. However, this function had a minimal effect on the output of the model (Fig. S4H) and was therefore not implemented in generating the final outputs shown. *In vivo* analysis of RNAPI stalling in the absence of topoisomerase activity indicated major stall sites at around +2Kb (El Hage et al., 2010). We postulate that this represents the point at which, due to the increasing size of the nascent pre-ribosome, the polymerase is unable to proceed further by rotation around the DNA (which does not generate torque), becoming fully torsionally entrained. We therefore included a function that progressively implements the effects of DNA torque - from 0 at the initiation site, where the polymerase can rotate ~freely around the DNA, to 100% at +2Kb (Fig. 4D). In the model this is the “Low Entrainment Region” (LER). We also introduced an elongation rate penalty over this region to account for the frictional cost of rotating the polymerase complex around the DNA. Importantly, within the LER neighboring RNAPI complexes can change relative positions without generating high torsional stress, potentially allowing more freedom to respond to effects of the nascent transcript.

##### 4. Effects of nascent transcript sequence

Folding of the nascent transcript was incorporated with high stability (low ΔG; calculated using a 65 nt of rolling window plus 15 nt off set) correlated with increased probability of rapid elongation and decreased probability of backtracking. The correlation between RNAP density and stability of the RNA:DNA duplex in the transcription bubble was incorporated with high stability (low ΔG; calculated using an 8 nt rolling window) correlated with decreased probability for rapid elongation. The effect of each feature was calculated for every nucleotide position. For ease of implementation these were combined in the model as “RNA Elements”.

The model was implemented in MATLAB and the full code is available as a git repository: https://bitbucket.org/bdgoddard/rnap_public/src/master/.

### Modelling indicates a major role for RNA folding

We constructed a set of dynamic models, which were run to achieve equilibrium states (Figs. 4B, C, E-G).

Discrete Stochastic Elongation alone generated a uniform distribution along the rDNA, since each polymerase moves independently with a stochastic distribution of step times and variability generated by stochastic initiation (Fig. 4B). A model implementing DNA torsion alone gives a broadly similar, relatively uniform profile (Fig. 4C). All polymerases are constrained to move as a single convoy, with DNA torsion effects between polymerases accelerating and periodically stalling elongation. Neither of these models closely match the *in vivo* EM and CRAC data. Inclusion of a 5’ Low Entrainment Region generated a distribution that more closely matched the *in vivo* data, since we now see a clear 5’ bias in modeled RNAPI density, with polymerases moving more slowly and more closely positioned over the initial 2Kb (Fig. 4E). The model including only the RNA Elements generated a highly uneven polymerase distribution, reflecting differences in folding energy and base composition across the entire rDNA (Fig. 4F). Finally, incorporating all of these features into a single model gave a distribution closely approximating the *in vivo* data (Fig. 4G and 4I). This shows both the 5’ enrichment and relatively discrete peaks observed in the EM and CRAC data. As a potential source of the 5’ bias we also considered premature termination of transcription, however this decreased number of RNAP per gene by 30% to achieve a similar 5’ bias (Figs. S4I and S4J) and therefore was excluded as a key factor in the model.

Alignment of peak and trough locations from the model with the experimentally derived peaks and troughs showed a clear overlap (Figs. 4H; modeled data in grey, CRAC data in green). This confirmed that the model significantly recapitulates the experimental data at high resolution. Major discrepancies are speculated to reflect sites at which backtracking is limited by stable binding of trans-acting factors rather than stem structures; see Discussion.

In the final model, the relative contribution of forces from different elements is clearly dominated by RNA folding (Fig. 4J), whereas DNA torsion has the weakest impact at each elongation step. However, entrainment alters elongation kinetics in the same direction over multiple steps to maintain relative RNAPI positions.

A striking conclusion from the model concerns the combined effects of the different features on the probability of RNAPI backtracking and collisions (Fig. 5). Stochastic elongation alone generates a low frequency of backtracking but a high frequency of collisions (Fig. 5A and B). Inclusion of torque, generated from DNA torsion, reverses this: increased probability of backtracking and reduced probability of collisions. Both backtracking and collisions are substantially suppressed by also including RNA structure (RNA Elements). The final model suggests that RNAPI takes advantage of both a low frequency of backtracking thanks to RNA structure and a low level of collisions due to DNA torsion.

**Figure 5.**
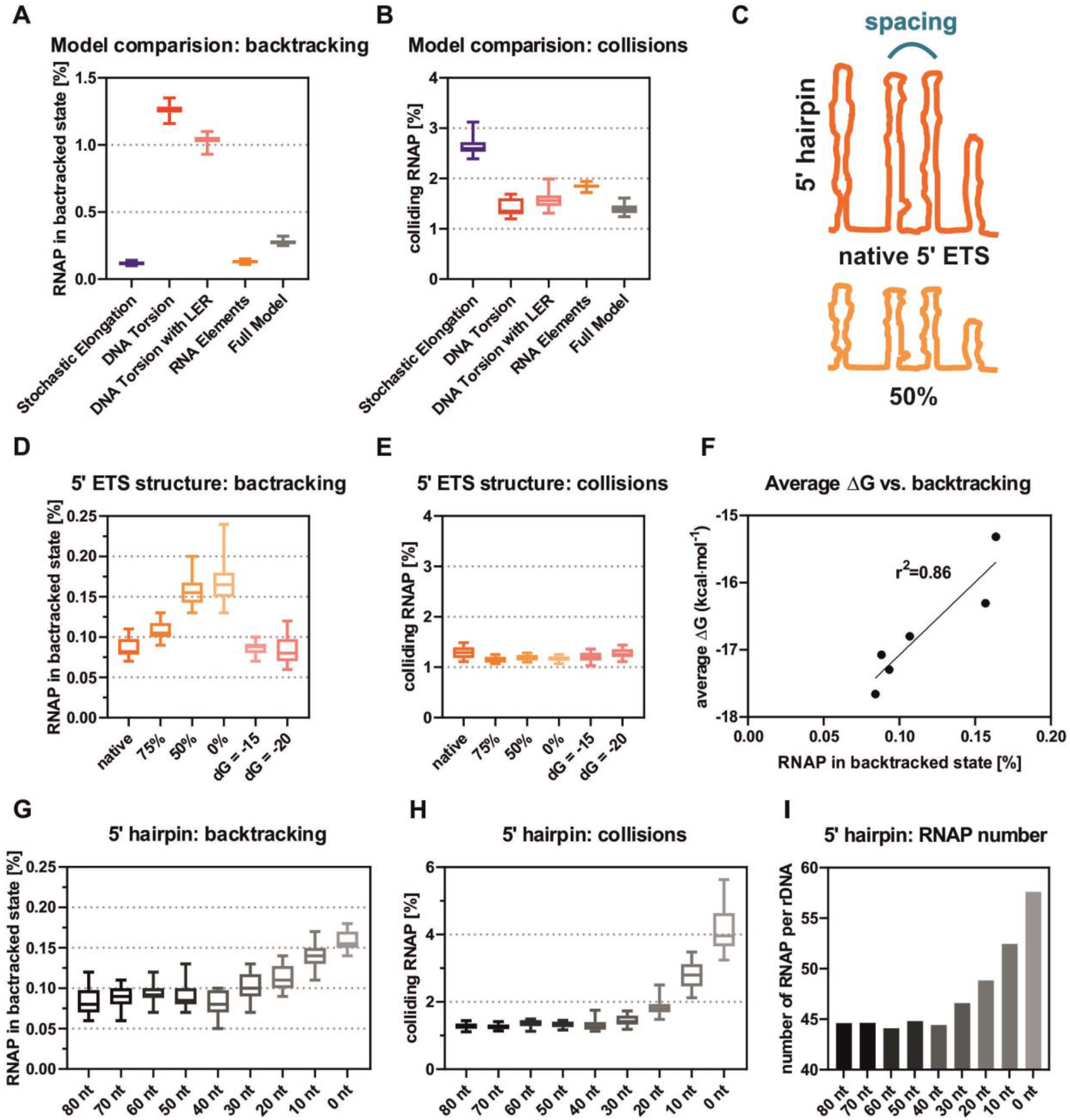
Conclusions from the model of RNAPI transcription. A: RNAP in backtracked state. Backtracking events were saved for each of the simulations (Fig 6B-F). Percentages of RNAP found in the backtracked state in different models were calculated and are presented as a boxplot. B: Frequency of colliding RNAP. Calculated as (A). C: Overview of the 5’ ETS structure modifications. Changing spacing between hairpins (top panel) or reducing strength of RNA structures (bottom panel). D: Frequency of RNAP backtracking in response to weaker 5’ ETS structures. The 5’ ETS with RNA structures reduced to 75%, 50%, 0% or fixed to ΔG = −15/−20 kcal·mol^−1^ was used. E: Frequency of colliding RNAP in response to weaker 5’ ETS structures. F: Frequency of RNAP in backtracked state is correlated with average stability of RNA structures. ΔG calculated for entire rDNA with modified 5’ ETS. G: Frequency of RNAP in backtracking in response to altered 5’ hairpin positioning within the 5’ ETS. The 5’ ETS with fixed ΔG = −20 kcal·mol^−1^ was used to test the impact of changes in 5’ structures. H: Frequency of colliding RNAP in response to the altered 5’ hairpin positioning. I: Number of RNAP per rDNA in response to the altered 5’ hairpin positioning.

The presence of a strongly folded 5’ ETS region in the pre-rRNA is conserved among eukaryotes. However, the primary sequence and length of the 5’ ETS are variable between species. We therefore assessed how overall folding of the 5’ ETS affects transcriptional output, by modeling a set of alternative structures (Fig. 5C); with 1) decreased ΔG over the 5’ ETS (Fig. 5A), or 2) altered spacing between the hairpins (Fig. 5C). For this analysis, the impact of the transcription bubble sequence was disregarded.

Consistent with results in Fig. 4, decreased folding energy of the 5’ ETS region caused increased RNAP backtracking (Fig. 5D), whereas collisions (Fig. 5E) and the total number of RNAP particles (Fig. S5B) remained unaffected. The fraction of backtracked RNAP correlated with the average ΔG over the 5’ ETS (Fig. 5F). Surprisingly, modification of spacing between the 5’ ETS hairpins (see overview in Fig. S5C) did not strongly affect output from the simulation (Fig. S5D-F). All three measures, backtracking, collisions and RNAP number, were very similar to the native 5’ ETS. Together these results indicate that strong secondary structures in the 5’ ETS are functionally important in reducing RNAP backtracking.

The 5’ proximal hairpin in the 5’ ETS seems to be unique. The RNAPI footprint determined by cryo-EM was estimated at 38 bp. Any strong structure appearing within this region would potentially accelerate promoter clearance. To test this, we generated a set of *in silico* constructs with the 5’ ETS fixed to dG = −20 kcal·mol^−1^ starting at positions 0 nt, 10 nt etc. - up to 80 nt. The 5’ ETS with structures starting at early positions (0-30 nt) showed increased backtracking and collisions (Figs. 5G and 5H), with increased rDNA loading (Fig. 5I) reflecting the effective initiation rate. Analysis of structures and folding energy of the authentic 5’ hairpin reveals very weak intermediate structures in comparison to the full-length hairpin (Fig. S5G). This feature is conserved in the human 5’ ETS (Fig. S5H) and in other species with characterized 5’ ETS sequences (*S. pombe* and *M. musculus*), all of which have very weak or absent 5’ terminal structures. We speculate that this lack of short, stable 5’ structures is a conserved feature that reduces overloading of the rDNA transcription unit.

### Effects of RNA folding are widespread and have regulatory potential

The key conclusions derived for RNAPI are expected to hold for all other polymerases and species. We therefore assessed the effects of nascent transcript structure for *Schizosaccharomyces pombe* RNAPI, and other RNA polymerases in budding yeast.

The *S. pombe* RNAPI CRAC profile revealed an uneven distribution with a 5’ bias (Fig. 6A), similar to *S. cerevisiae* RNAPI (Fig 1D). Metaplot analysis of RNAPI density troughs, versus folding energy of the nascent transcript, revealed a strong correlation (Fig. 6B, p=0.005).

**Figure 6.**
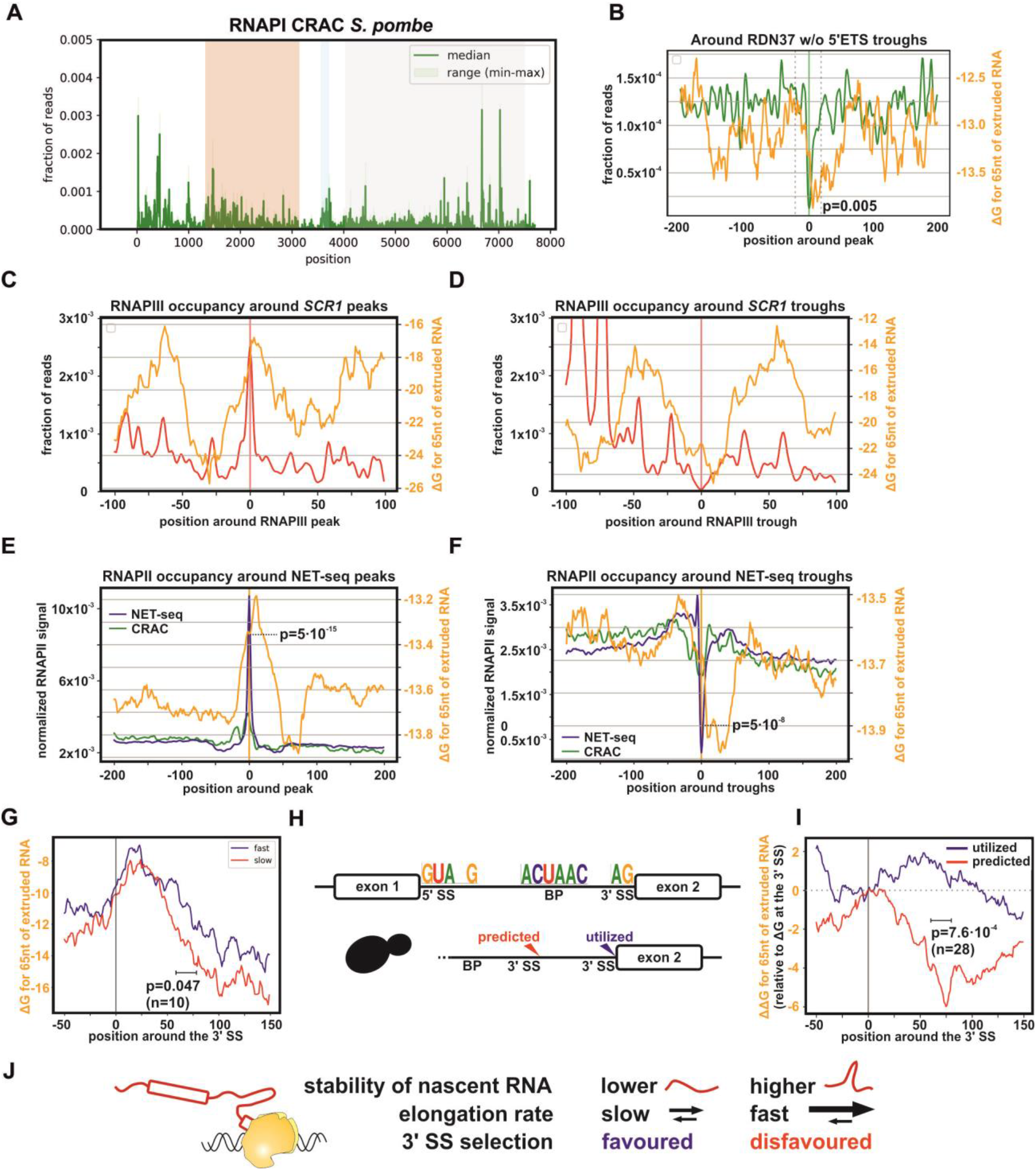
Effects of transcript folding are widespread and have regulatory potential. A: *S. pombe* RNAPI CRAC distribution over the gene encoding the pre-rRNA. Two biological replicates. Annotation as Fig. 1D. B: *S. pombe* RNAPI CRAC trough metaplot for entire pre-rRNA with folding energy of the nascent transcript. C: RNAPIII CRAC peak metaplot compared to predicted folding of nascent scR1 ncRNA. Folding energy (ΔG in Kcal mol^−1^) was calculated for a rolling 65 nt window behind RNAPIII, offset by 15 nt. D: As D except that CRAC troughs were compared to folding of nascent scR1. E: Metaplot showing peaks of NET-seq density for RNAPII. Peaks from the top 50% of RNAPII transcripts longer than 300 nt were overlaid with the folding energy of the nascent transcripts. Folding energy (ΔG in Kcal mol^−1^) was calculated for a rolling 65 nt window behind RNAPII, offset by 15 nt. F: As E but showing troughs in NET-seq density. G: Folding of nascent RNA around the 3’ SS of pre-mRNA genes classified as fastest third and slowest third for spliced, non-ribosomal protein genes (Barrass et al., 2015). H: Conserved feature of yeast introns used for *de novo* prediction (top panel). Genes containing a predicted but skipped 3’ SS were analyzed (n=28, bottom panel). I: Relative folding energy of nascent RNA around utilized 3’ SS, versus predicted but skipped 3’ SS, relative to the 3’ SS (position 0). J: Suggested role of stability of nascent RNA in selection of the 3’SS.

RNAPIII generally transcribes very short pre-tRNA transcript. However, the RNAPIII-transcribed *SCR1* gene encodes the 522 nt long scR1 ncRNA component of signal recognition particle. Previous RNAPIII CRAC data showed a very uneven distribution across *SCR1* (Turowski et al., 2016). A peak and trough metaplot for RNAPIII density, versus the folding energy of the nascent scR1 RNA, revealed a high degree of correlation (Figs. 6C-6D and S6A), similar to that observed for RNAPI, However, the number of features was too low to perform statistical analysis.

Published, high-resolution analyses of RNAPII distribution by either NET-seq or CRAC, using the catalytic subunit Rbp1, also reveal strikingly uneven density (Churchman and Weissman, 2011; Milligan et al., 2016) (Fig. S6B). Independent biological replicates for Rpb1 distribution in NET-seq and CRAC showed good reproducibility across well-transcribed genes (Fig. S6C), indicating that the fluctuations represent genuine differences in RNAPII density.

Some of the variation in RNAPII occupancy reflects nucleosome positioning, with maximal density (minimal RNAPII elongation rate) seen at the center of nucleosomes (Churchman and Weissman, 2011; Milligan et al., 2016), which are generally well-positioned in yeast. To determine whether structure in nascent transcripts also affects RNAPII occupancy, we used a peak-calling algorithm to define peaks and troughs in the RNAPII density, across 50% of the most highly transcribed genes that are longer than 300 nt (n=1073). We used NET-seq peaks to generate metaplots, since published RNAPII CRAC data were prepared using a protocol that does not specifically recover the nascent 3’ end. This showed a correlation between the RNAPII peaks (n=9927) and troughs (n=4776) and the rolling average of predicted ΔG (shown for a 65 nt window with 15 nt offset in Figs. 6E and 6F). RNAPII occupancy peaks were associated with a clear peak of folding energy, whereas troughs, indicating rapid elongation, were associated with stronger nascent RNA structure (Fig. 6E, p=10^−7^ T-test and p=5·10^−16^ Wilcoxon signed-rank test, and 6F, p=7·10^−6^ T-test and p=5·10^−8^ Wilcoxon signed-rank test, orange line).

To determine whether nascent RNA structure may have a regulatory potential, we used pre-mRNA splicing as a model process. The splicing machinery co-transcriptionally recognizes the 5’ splice site (SS), branch point (BP) and the 3’ SS (acceptor site). A prediction derived from our findings is that stronger structure in the nascent RNA should reduce the time available for co-transcriptional selection of the 3’ SS, disfavoring rapid, cotranscriptional splicing. Analyses using extremely fast metabolic labeling, previously ranked yeast pre-mRNAs by splicing speed (Barrass et al., 2015) (Fig. S6D). Consistent with our prediction, the fastest third of spliced genes had less structure in the nascent RNA at the start of exon 2 when compared to the slowest third of spliced genes (Fig. 6G, p=0.047 for n=10).

The 3’ SS consensus is notably weak, consisting of only two bases (AG) which are expected to co-occur frequently. This suggests a kinetic model for 3’ SS selection, based on a “window of opportunity”. To assess the potential role of nascent RNA folding as a decisive factor, we defined all yeast introns *de novo* using previously described features (Fig. 6H) and specifically focused on those which possess a predicted, but unutilized, 3’ SS upstream of the authentic site. Then we compared changes in folding energy of the nascent RNA downstream from the predicted and utilized 3’ SS (Fig. 6I for ΔΔG, p=7.6·10^−4^ and S6E for ΔG, p>0.05). Nascent RNA extruded after transcription of the utilized 3’ SS maintained RNA folding (ΔΔG) on similar level whereas predicted but unutilized 3’ SS RNA are accompanied by stronger folding of nascent RNA (ΔΔG). Interestingly, this would suggest that relative folding energy (ΔΔG) is more important for selection of the 3’ SS, since overall stability of nascent RNA was not significantly different (Fig. S6E, p>0.05). According to our model, stronger folding of the nascent RNA speeds up RNAPII and decreases the “window of opportunity” for splicing to occur. We propose that this favors skipping of the unused, potential 3’SS (Fig. 6J). Notably, significant differences were seen for folding energy nascent RNA, even when the folding window did not include the 3’SS, making it unlikely that direct effects on the structure or accessibility of the acceptor site are responsible for the observed correlations.

We conclude that stimulation of transcription elongation by nascent RNA structure is a conserved feature of all three eukaryotic polymerases and regulation of co-transcriptional processes is, at least partially, determined by local folding of nascent RNA.

## DISCUSSION

The process of transcription is fundamental to all gene expression. Structures of RNA polymerases have been determined at high resolution from many systems and the mechanism of nucleotide addition is characterized in considerable detail. However, the kinetics of initiation and elongation *in vivo* are less well understood. Here we aimed to better understand RNA transcription elongation, by providing a bridge between *in vitro* biochemical experiments and *in vivo* biology, and between the mechano-physical properties of polymerases and genome wide analyses.

Analyses of eukaryotic transcription by multiple techniques have revealed uneven polymerase occupancy, reflecting variable elongation rates. This is important, because many RNA processing factors act very quickly on the nascent transcript. For example, splicing of pre-mRNA is strikingly speedy in yeast (Wallace and Beggs, 2017), although more heterogeneous in metazoans (Alpert et al., 2017; Drexler et al., 2019), and this can shape the mature mRNA through alternative splicing (Saldi et al., 2016). Understanding the detailed kinetics of transcription elongation *in vivo* will therefore be predictive for processing decisions.

In eukaryotes, RNAPI is most amenable to these analyses, since it transcribes only a single product from the nucleosome-free rDNA. Our EM analyses of Millar chromatin spreads indicated that the distribution of RNAPI was uneven across the rDNA transcription units, with an excess of polymerases in the 5’ region. In an orthogonal approach, we determined the distribution of RNAPI by CRAC UV crosslinking. This confirmed the 5’ enrichment for RNAPI density, but also revealed a strikingly uneven, local polymerase distribution, most notably over the 5’ ETS region of the pre-rRNA (Fig. 1D).

Analysis of features that correlate with peaks and troughs of RNAPI density showed a modest correlation with the stability of the RNA-DNA duplex in the transcription bubble, but strong correlation with the calculated folding energy of the nascent pre-rRNA transcript close to the polymerase (Fig. 2). RNA polymerases operate as Brownian ratchets and are prone to backtracking, which serves as a proofreading step (Fig. 1A). During backtracking, the newly transcribed RNA must re-enter the polymerase. However, the channel accepts only single stranded RNA, suggesting that backtracking will be resisted by any RNA structures that form sufficiently rapidly in the nascent transcript. RNA hairpins were previously suggested to stabilize pauses or prevent backtracking of bacterial RNAP (Dangkulwanich et al., 2014).

Using RNAPI transcription *in vitro*, we confirmed that strong structures in the nascent transcript effectively resist backtracking and defined the stability of stems that can either block or slow backtracking by RNAPI (Fig. 3). Additionally, our genome wide data provide evidence that RNA structure substantially modulates transcription elongation by RNAPII and RNAPIII.

Since only naked, single stranded RNA can re-enter the polymerase exit channel, we predict that trans-acting factors with sufficiently rapid and stable bonding to the nascent RNA will also resist backtracking. Supporting this conjecture, we note that there were fewer discrete 5’ ETS peaks in the model than in the CRAC data. The prominent CRAC peak around +100 corresponds with the major binding site for the UTP-A complex of early-binding ribosome synthesis factors (Hunziker et al., 2016; Sun et al., 2017), which were previously implicated in transcription and designated t-Utps (Gallagher et al., 2004). Similarly, a CRAC peak further 3’ is close to the major U3 snoRNA binding site at +470. We postulate that RNA packaging factors bound to nascent transcripts also function as ratchets, favoring progressive RNAPI elongation.

To better understand the contributions of different features to the behavior of RNAPI *in vivo*, we developed a mathematical model of rDNA transcription. Multiple simulations were run and compared to the RNAPI distribution determined by *in vivo* analyses (Fig. 4 and 5). Basic parameters were taken from published literature (as described in Extended Text), defining the initiation frequency, overall elongation rates, the torque generated by torsion in the DNA and the time distribution of the stochastic elongation and backtracking steps. Effects of DNA torsion were previously included in models for RNAPII convoys visualized following promoter bursting (Lesne et al., 2018; Tantale et al., 2016) and the cooperative behavior of bacterial polymerases (Heberling et al., 2016; Kim et al., 2019). Other features were fitted to the experimental data, including the negative correlation of elongation rate with high G+C content in the transcription bubble and the positive effects of stable folding of the nascent transcript. A major criterion of the model optimization was a high yield of transcriptional output, since this is a crucial feature of RNAPI transcription (Fig. S4F and S4J, see Extended text for details).

The graphical summary of the model is presented in graphical abstract. A notable finding from the model was that the inclusion of the effects of torque reduces the numbers of colliding RNAPI but increases the fraction of RNAPI in a backtracked position. Addition of the effects of nascent transcript folding reduced the frequency of backtracking, while retaining the low level of collisions. This underlined the positive contribution of RNA structure to productive elongation.

Additional problems with RNAPI transcription may occur *in vivo*; e.g. deep backtracking, prolonged pausing leading to premature termination or very long r-loops. However, we predict that these severely disrupt transcription on the affected rDNA unit and are not considered in the model.

The yeast 5’ ETS is notably highly structured, particularly as it has a low G+C content in keeping with the overall genome. The overall ΔG for the 700 nt ETS is around −265 Kcal mol^−1^, whereas the adjacent 700 nt of the 18S rRNA has a ΔG of −220 Kcal mol^−1^, despite the well-known tendency for rRNAs to be highly structured. We therefore speculated that the high degree of 5’ ETS structure at least partly reflects selection for structures that promote efficient transcription. Supporting this assumption, we found that structures within the 5’ ETS decreased the frequency of RNAPI backtracking, and this effect correlated with the overall ΔG of the nascent RNAs. Overall, we postulate that strong structures within the transcribed spacers reduce backtracking of RNAPI. However, evolution of the exact structure and sequence was driven by the processing machinery rather than efficiency of transcription. Notably, relatively sharp peaks of RNAPI density correlated with the apexes of the extended stem structures in the ETS. We speculate that this arises because the lowest enhancement of elongation resulting from RNA structure occurs at these sites. Weaker, transient structures will have formed during extrusion of the 5’ sides of the extended stems, giving some boost to elongation, but these must be unfolded prior to refolding into the extended final stems.

Our key findings on the effects of folding in the nascent transcript on polymerase elongation, are also applicable to RNAPI from *S. pombe* and to RNAPII and RNAPIII from *S. cerevisiae* – and presumably to RNA polymerases in many or all other systems. While folding energy of the nascent transcript emerged as a the most significant feature in determining RNAPI elongation rates, its role in RNAPII elongation is expected to be tempered by the many other factors impacting on elongation, notably nucleosome structure, polymerase modification and association with cis-acting elongation factors. Despite this, the correlation between RNA folding and polymerase density was clearly seen, both by CRAC crosslinking and in NETseq data, which use orthogonal approaches. We note that the average change in folding energy for RNAPII transcript is lower than for the rDNA 5’ ETS (Fig. 6E vs. 2D), but the correlations are supported by very significant p-values (Fig. 6). RNAPII data are affected by presence of nucleosomes and lower sequencing depth (the signal is divided over many genes), affecting application of the peak-calling algorithm. Moreover, the folding window was optimized for the hairpins within the pre-rRNA 5’ ETS. The results could be improved by calculating the change in cotranscriptional folding as each nucleotide is extruded, but this is not currently technically feasible. Measurements of the effects of nascent RNA on RNAPII dynamics *in vitro* using single-molecule approaches (Zamft et al., 2012) support our findings *in vivo*. We conclude that RNAPII dynamics are substantially influenced by folding of the nascent transcript *in vivo*, despite many additional factors.

It is a longstanding observation that signals within pre-mRNAs defining splice sites have surprisingly little information content, relative to the specificity of splicing. Multiple additional features contribute to accurate splice site selection but, to the best of our knowledge, these were not previously reported to include the nascent structure of RNA close to the polymerase. The relationship between RNA structure and splicing efficiency in yeast was previously assessed, but focusing on intronic sequences that were assumed to be critical (Barrass et al., 2015). In the yeast data, RNA folding at the 3’ splice site is unlikely to be responsible for the effects on splice site selection. Rather, we propose that unstructured RNA downstream of the intron favors slowed elongation, which is associated with rapid splicing and proximal 3’ splice site selection. In contrast, structured RNA promotes rapid elongation, which is associated with relatively slow splicing and distal splice site usage. We speculate that slower elongation of RNAPII facilitates splicing by allowing more time for recognition of the 3’ SS by splicing factors associated with C-terminal domain of the polymerase. Notably, the “window of opportunity” for 3’ SS recognition is presumably substantially shorter than the actual pre-mRNA splicing reaction that is assessed by transcript sequencing in yeast and mammalian systems (Alpert et al., 2017; Drexler et al., 2019; Neugebauer, 2019; Wachutka et al., 2019; Wallace and Beggs, 2017). Finally, we note that similar considerations potentially apply to other cotranscriptional events that depend on RNAPII-associated recognition, including alternative polyadenylation.

## Supporting information

Supplemenal File Combined

## Acknowledgements

We thank Grzegorz Kudla for critical reading of the MS and the members of Tollervey group and Guido Sanguinetti for stimulating discussions. We thank Christoph Engel, Joachim Griesenbeck and Michael Pilsl for support and valuable advice in setting up the *in vitro* assay. We thank Tomas Gedeon, Lisa David and Tamra Heberling for stimulating discussions and valuable advice about the mathematical model. This work was funded by Wellcome through a Principle Research Fellowship to D.T. (077248). TWT was supported by the Polish Ministry of Science and Higher Education Mobility Plus program (1069/MOB/2013/0). SLF was supported by the NIH (GM06952). Work in the Wellcome Centre for Cell Biology is supported by a Centre Core grant (203149).

## Data access

All RNA sequencing data from this study have been submitted to the NCBI Gene Expression Omnibus (GEO; http://www.ncbi.nlm.nih.gov/geo/) and are available under accession number GSE136056. The full MATLAB code for the mathematical model has been submitted as a git repository: https://bitbucket.org/bdgoddard/rnap_public/src/master/.

