## Supplementary material for "Nascent transcript folding plays a major role in determining RNA polymerase elongation rates": Supplemenal File Combined

**Supplementary Materials**

**Turowski et. al**

**Extended Text**

Table S1.

Table S2.

**Supplementary Figures**

Fig. S1

Fig. S2

Fig. S3

Fig. S4

Fig. S5

Fig. S6

**Supplementary Tables**

Table S3.

Table S4.

Table S5.

**Materials and methods**

### Extended Text

#### Validation of RNAPI CRAC data

Two major aspects of the CRAC data were investigated: Contamination with mature rRNA or processed pre-rRNA and bias in sequence recovery.

In total RNA, mature rRNAs (18S, 5.8S and 25S rRNA) are much more abundant than the spacer regions (5'ETS, ITS1, ITS2, 3'ETS) present in the nascent transcript. However, the recovery of reads mapping to the rRNA sequences was not clearly elevated relative to the spacers and there was no accumulation at the mature rRNA boundaries (Fig. 1D). This shows that the RNAPI CRAC data are not significantly contaminated by mature rRNAs.

During pre-rRNA transcription, the nascent transcript is cleaved at four sites; A0, A1, A2 and B0. Cleavages at A0-A2 are coupled and predominately cleaved in the nascent transcript, but processing occurs when RNAPI has travelled ~1.2Kb downstream from site A2 (Axt et al., 2014; Kos and Tollervey, 2010). Sequences terminating at sites A0, A1 and A2 were not elevated in the CRAC data (Fig. 1D and S1C), confirming that the processed pre-rRNAs were not recovered. Mapped 3' ends from cDNAs are therefore expected to represent the positions of *bona fide* 3' ends in nascent transcripts.

We also performed experiments to validate the Rpa190 CRAC data and detect potential bias in target recovery. Notably, all of these analyses yielded RNAPI distributions that were consistent with the results of CRAC with Rpa190 (Fig. 1F).

1: To reduce the possibility of non-specific cross-linking to Rpa190, we analyzed a range of shorter UVC cross-linking times. These showed minimal changes (Fig. S1D).

2: To exclude steric preferences in RNA cross-linking, we HTP-tagged the second largest subunit of RNAPI, Rpa135 (Fig. S1E). This showed a similar 5' bias to Rpa190, and substantial overlap at the level of individual peaks, as shown by a peak metaplot (Fig. S1E, embedded panel; see Fig. S1F and Material and Methods for details on peak metaplot generation).

3: We performed a PAR-CRAC experiment, in which RNA was metabolically labeled with 4-thiouracil (4SU) and cross-linked using UVA (Fig. S1G). 4SU crosslinking involves different photochemistry and may be less prone to recover non-specific crosslinking relative to UVC (Shchepachev et al., 2019). The peak metaplot for PAR-CRAC was very similar to the CRAC data (Fig S1G). However, some enrichment for U-rich sites was observed in PAR-CRAC, as expected (Fig. S1H).

4: Wild-type yeast strains generally have ~150-200 ribosomal repeats, of which around 50% are reported to be actively transcribed, making it conceivable that the apparent 5'-end bias (Fig. 1D) arises from premature termination on "inactive" repeats. To test this possibility, Rpa190 CRAC was performed in a yeast strain with only 25 rDNA repeats, all of which are highly transcribed. The RNAPI profile in this strain was almost identical to the wild-type (Fig. S1I).

5: We considered the possibility that RNA interacting with the outside of the polymerase might contribute to the signals, although the requirement that recovered RNA has a 3' OH group made this unlikely. To test this, we considered only cDNA sequences shorter than 20 nt, since this region will be almost entirely located within the transcription bubble and RNA exit channel. This analysis also revealed the distinctive peaks for RNAPI distribution (Fig. S1J).

6: Finally, we considered bias originating from the CRAC experimental protocol. Mainly the relationship between nascent RNA recovery, RNA structure, UV crosslinking and adapter ligation steps during library preparations. The arguments against this hypothesis are as follows: (1) nascent RNA interacting with RNAP is buried inside the channel in its extended, unstructured form, therefore, there should be no influence of structure on the UV crosslinking efficiency. (2) The CRAC protocol involves highly denaturing conditions to reduce the background. Following protein denaturation, the RNA could be susceptible to folding, potentially sequestering RNA ends and hindering adapter ligation. In such a case we would expect lower recovery of highly structured RNAs, e.g. hairpin loop regions. However, this is in marked contrast to our results (Fig. S2F).

From this validation we conclude that CRAC approximates the genuine distribution of RNAPI at most sites along the rDNA transcription unit *in vivo*. All subsequent analysis was performed using the median of six biological replicates (Fig. S1K). Moreover, we generated randomized datasets and compared RNAPI CRAC with them using a Spearman test (Fig. S1L). This revealed that RNAPI CRAC data present a non-random distribution.

**Mathematical model of RNAPI transcription**

The numerical model for elongation steps of RNAPI transcription kinetics, was developed using input data taken from biological experiments wherever possible (Table S1).

**Table S1.** Fixed parameters used in the mathematical model of RNAPI transcription.

| Name | value | Reference |
|---|---|---|
| Mean elongation of RNAPI | 40 nt $s^{-1}$ | (Kos and Tollervey, 2010) |
| 35S pre-rRNA transcription time | 170 s | (Kos and Tollervey, 2010) |
| rDNA repeats | 150-200 | (Nomura, 1999) |
| Active rDNA repeats | 75-100 | (Toussaint et al., 2005) |
| RNAPI per rDNA | 50±2.5 | (Dasgupta et al., 2007; El Hage et al., 2010; French et al., 2003; Hontz et al., 2008; Oakes et al., 2006; Sandmeier |

| | | |
|---|---|---|
| | | et al., 2002; Schneider et al., 2006; Tongaonkar et al., 2005; Viktorovskaya and Schneider, 2015) |
| 35S rDNA length | ~7000 | SGD |
| RNAPI footprint | 38 nt | (Neyer et al., 2016) PDB 5M5X |

*Justification of parameters of the model*

1. Quantification of molecules of RNA polymerases

To estimate total copy numbers for RNAPI, RNAPII and RNAPIII, we re-analyzed three independent studies: (Chong et al., 2015; Ghaemmaghami et al., 2003; Kulak et al., 2014). An average and median for all subunits were calculated for each RNA polymerase (Fig. S4A). These calculations were repeated for all specific subunits for each RNA polymerase and presented similar trend. Data expressed in arbitrary units (Chong et al., 2015) were used only to confirm ratios between RNA polymerases. Analysis of these data indicated that RNAPI and II are present at similar levels of 5,000 - 6,000 molecules per cell, whereas RNAPIII is present in 2,500-3,000 copies.

2. Transcription initiation rate

Rapidly dividing yeast cells produce ~200,000 ribosomes per generation (~100 min), corresponding to ~2,000 ribosomes $min^{-1}$. These are transcribed from 75-100 rDNA repeats. Each transcription unit should therefore release ~20-27 completed pre-rRNA transcripts per minute (1 transcript every 2.2 - 3 sec). The transcription initiation rate cannot therefore be less than 1 initiation per 2.2 - 3 sec, but might be greater if the processivity of RNAPI is less than 100% or the elongation rate is non-uniform.

Transcription initiation by RNAPI has undoubtedly evolved to be extremely efficient. We postulate that polymerase may be recruited to the rDNA promoter faster than the time needed for the previous polymerase to clear the initiation site, making promoter clearance rate limiting. Therefore, we modeled RNAPI transcription initiation as a stochastic process with a success probability between 0.33 and 1 per second.

3. RNAPI number per rDNA transcription unit and RNAPI spacing.

The maximum average number of RNAPI molecules per rDNA transcription unit can be estimated from the number of RNAPI in the cell (5,000-6,000 molecules) and rDNA repeats (75-

100), giving a range of 50-60. This figure is in good agreement with quantification of RNAPI complexes from Miller chromatin spreads (~50, Table S1). The number of RNAPI molecules on the 7 Kb long rDNA transcription unit gives an average RNAPI spacing of 120-140 nt.

$$spacing = \frac{7000\,nt}{\left(\frac{RNAPI\ molecules}{active\ rDNA\ repeats}\right)}$$

This value is in good agreement to independent calculations derived from metabolic labelling experiment (Kos and Tollervey, 2010). The average velocity of RNAPI (40 nt $sec^{-1}$) and transcript release rate (1 per 3 sec; from initial calculations above) predicts a spacing of 120 bp. Measurements of the relative positions of RNAPI in Miller spreads by tomography, indicated minimal center to center separation of 15 nm (Neyer et al., 2016), which is estimated to reflect a 44 bp. This figure may therefore represent a minimal spacing between RNAPI molecules *in vivo*.

4. Elongation rate of RNAP in the discrete model

The velocities of RNA polymerases have been determined *in vivo* and *in vitro* many times and some examples were summarized in Table S2. Interestingly *in vitro* measurements are systematically lower than *in vivo*.

**Table S2.** The velocity of RNA polymerases

| RNA polymerase | Method | Elongation rate nt $sec^{-1}$ | Reference |
|---|---|---|---|
| RNAPII human | single cell microscopy, *in vivo* | 115 | (Tantale et al., 2016) |
| RNAPII yeast | single molecule transcription assay, *in vitro* | 25-30 | (Dangkulwanich et al., 2013) |
| RNAPII yeast | single molecule transcription assay, *in vitro* | 18.7 +/- 2.7 | (Lisica et al., 2016) |
| RNAPI yeast | single molecule transcription assay, *in vitro* | 32.2 +/- 2.5 | (Lisica et al., 2016) |
| RNAPI yeast | Metabolic labelling, *in vivo* | 40 | (Kos and Tollervey, 2010) |
| RNAPII human | 4sUDRB-seq, *in vivo* | 30-100 | (Fuchs et al., 2014) |

| | | | |
|---|---|---|---|
| RNAPII human | DRB-arrest release, *in vivo* | 50 | (Cortazar et al., 2019) |

Approximate RNAPI elongation rates can be also obtained from number of ribosomes produced per generation (200,000), yeast doubling time (100 min), pre-rRNA length (~7000 nt) and number of transcribing RNAPI molecules (5000 - 6000).

$$V_{RNAPI} = \frac{200\,000\,ribosomes\ \cdot 7000\,nt}{6000\,seconds\ \cdot RNAPI\ molecules}$$

Based on the published data, ~40 nt·sec$^{-1}$ is expected to be the overall average velocity of transcribing RNAPI. However, pause-free elongation is very unlikely *in vivo*. Therefore, we used 50 nt sec$^{-1}$ as the intrinsic, average RNAPI elongation rate ($V_{Int}$) in our model.

The *in vitro* velocity distribution for RNA polymerase from *E. coli* was determined using single-molecule measurements (Adelman et al., 2002). It is described by two Gaussian functions: The first function, comprising 7.8% of the area, represents the paused state and is centered at 0.9 nt sec$^{-1}$ (Fig. S4B, red line). The second function reflects active elongation and centered at 12.8 nt sec$^{-1}$ (Fig S4B, green line). We adopted this function for RNAPI, using an *in vivo* elongation velocity centered at 50 nt sec$^{-1}$ (Fig. S4B').

RNA polymerase elongation is based on a Brownian ratchet mechanism, in which each step of elongation and catalysis is discrete and independent from other steps. Classical mechanics and momentum do not apply to molecular processes and we therefore constructed a stochastic and discrete model. At each time step, there is a probability of moving 0, +1 or -1 nucleotides according to the distribution presented in Fig. S4B`.

$V_{Elongation} = N(V_{Int}, (\sigma \cdot V_{Int})^2)$, where $V_{Int} = 50$ and $\sigma = 0.4$

$V_{Paused} = N(V_{Paused}, \sigma_{Paused}{}^2)$, where $V_{Paused} = 0.9$ and $\sigma_{Paused} = 1.5$

For each particle, with probability $p = 0.078$, $V_{Random}$ is drawn from the $V_{paused}$ distribution, and otherwise from the $V_{elongation}$distribution.

Having computed the velocity $V_{Random}$ for a given time step of length $dt$, the corresponding probability of jumping in that time step is given by $p = |V| \cdot dt$. This is essentially the expected distance moved in one time-step. With probability $p$, the RNAP jumps in the direction of $V$ in that time step.

This probability can be modified by following factors:

(a) DNA torsion

(b) Promotion of RNAP elongation by nascent structure forming behind the polymerase

(c) Decrease of RNAP elongation by a strong RNA:DNA hybrid within the transcription bubble.

5. RNAP convoys imply DNA torsion effects

RNAP elongation along a DNA helix requires two types of movement: forward and rotary. In principal, either the DNA or polymerase can rotate with a frequency of ~240 rpm. The rDNA is nucleosome-free and loaded with multiple RNAPI complexes (~50 at 0.5 MDa each), each associated with up to 7 Kb of pre-rRNA transcript (up to 2.3 MDa) and a multi-megadalton pre-ribosome (6 MDa for the SSU processome alone) containing many assembly factors (Turowski and Tollervey, 2015).

The difficulty of moving these very large complexes through the highly viscous nucleolus environment (Bormuth et al., 2009), and steric problems that would be entailed by rapid rotation of the pre-rRNA around the DNA, make it very likely that the rDNA is rotated through an array of polymerases. If a group of RNAPI complexes move along the DNA together, this will not result in over- or under-winding of the DNA. This suggests that the RNAPI array on the rDNA acts cooperatively to rotate the DNA template. It is, of course, expected that DNA supercoiling (both, positive and negative) will be also alleviated by DNA topoisomerases I and II (Top1 and Top2). However, the abundance of Top1 is estimated to be very much lower than the RNA polymerases. Quantification reported by SGD (https://www.yeastgenome.org) based on multiple analyses: Top1; 4130 ± 2517 molecules per cell. Sum of the largest subunits of all three RNA polymerases; 33674 ± 12715.

We therefore propose that RNAPI complexes move as a group along single rDNA transcription unit while DNA rotates through the polymerases. Notably, similar models have been proposed for highly transcribed RNAPII genes associated with “convoys” of RNA polymerases resulting from transcriptional bursting (Lesne et al., 2018; Tantale et al., 2016), and for bacterial polymerase (Heberling et al., 2016; Kim et al., 2019). Finally, DNA rotation during transcription was observed directly in *E. coli* RNAP (Harada et al., 2001).

In the model for RNAP convoys, the distance between initially loaded RNAP molecules is maintained by torsion in the DNA helix. Transcription elongation of RNAP includes a translocation step based on Brownian motion. Only this step is assumed to be force sensitive (Dangkulwanich et al., 2013). Single-molecule elongation of bacterial RNAP can be stopped *in vitro* by application of a stalling force of 15-25 pN (Bustamante et al., 2004).

Overwinding and underwinding of DNA ($\sigma$) generates force. $\sigma = 0.00$ (0%) when DNA is relaxed (1 turn / 10.5 bp, 10 turns / 105 bp) and $\sigma = 0.10$ (10%) when DNA is 10% overwound (1.1 turn / 10.5 bp, 11 turns / 105 bp). When all polymerases within the convoy are moving along DNA with the same velocity (relative velocity $v_{rel} = 0$) the force generated by DNA torsion equals 0. However, when one polymerase moves faster than its neighbors ($v_{rel} > 0$), this results in DNA overwinding in front of RNAP and underwinding behind it (Fig. 4A). Both of these effects will favor slowing of the middle RNAP.

A DNA torque $\tau = 11\, nm \cdot pN$ was reported to stall bacterial RNAP *in vitro* (Ma et al., 2013). [Note that DNA torque and stalling force have different units.] An elegant solution was proposed to calculate a relationship between DNA torque $\tau$ and DNA overwind $\sigma$ (Fig. S4C) (Heberling et al., 2016).

$$\tau = \frac{\mu \pi^2 r^4}{10.5}\left[\ln(\frac{xP_{Init}}{dxP}) - \ln(\frac{xM_{Init}}{dxM})\right],$$

Where $\mu = 300\, pN/nm^2$ is the shear modulus for DNA, $r = 1\, nm$ is the radius of DNA, 10.5 is number of bases per turn. $\sigma$ is relationship between loading distance $xP_{Init}$ or $xM_{Init}$ and current distance $dxP$ or $dxM$. The current distance is calculated as $dxP = x_{n+1} - x_n$ and $dxM = x_n - x_{n-1}$ (Fig. 4A).

The force $F$ acting on the RNAP is calculated from DNA torque $\tau$ as previously described (Ma et al., 2013) (Fig S4D).

$F = \frac{\tau \cdot \theta}{d}$, where $\theta$ represents angular rotation of RNAP after 1 bp translocation 0.6 radian or 34°, converted from 10.5 bp per turn. $d$ is the contour length of DNA per bp (~0.34 nm).

It is notable that the value of sigma $\sigma$ causing RNAP stalling will be higher *in vivo* due to following reasons: (1) We assumed that highly packed and viscous environment of the nucleolus causes RNAPI to transcribe as a convoy. However, *in vivo* the ability of RNAPI to rotate around the rDNA will be greater than zero. Therefore, an increased limit of sigma $\sigma$ includes this capacity to spin around the rDNA without introducing an additional parameter. (2) The average velocities of bacterial RNAP or RNAPI *in vivo* are ≥ 2 fold higher than *in vitro* (Table S2). The previously developed function describing bacterial RNAP velocity in relation to DNA torque $\tau$ is based on *in vitro* data, hence we assume that RNAPI stalling force *in vivo* is higher and decided to increase sigma $\sigma$ appropriately.

We therefore used $\sigma = \pm 0.05$ as a parameter in our model. Much higher values are unlikely since $\sigma = \pm 0.20$ can lead to phase transition (Sarkar et al., 2001) and very low negative torque $\tau$ may lead to DNA melting.

In the model, DNA torsion modifies $V_{Random}$ as follows:

$$V_{Random+Torque} = V_{Random} + c\left(1 - \frac{xP_{Init} - b}{dxP - b}\right) - c\left(1 - \frac{xM_{Init} - b}{dxM - b}\right)$$

Where $xP_{Init}$ or $xM_{Init}$ are initial distances between polymerases when initiated (engaged on the DNA), $dxP$ or $dxM$ are current distances between polymerases, $b$ is the length of the transcription bubble (11 nt for RNAPI, PDB: 5M5X (Tafur et al., 2016)) and $c$ is a constant describing DNA stiffness. In the basic model for RNAP convoys, the initial separation of RNAP is established by the initiation rate, and then maintained by DNA torque.

6. Range of DNA stiffness constant c

The $V_{Random+Torque}$ equation allows calculation of the DNA stiffness constant $c$ in relation to DNA overwind $\sigma$. We use a simplified system of three RNAPs with an initial separation of 100 nt (as Fig. 5A). Then we solved the equation, in which a given value for $c$ should be strong enough to stop RNAP transcribing with average velocity $V_{Int}$ when % of DNA overwind $\sigma$ is equal to $r$:

$$c = \frac{-V_{Int}}{\frac{100}{100 + r} - \frac{100}{100 - r}}$$

$V_{Int}$ is the intrinsic velocity of RNAPI and equals 50 nt·sec$^{-1}$. This gives values of DNA stiffness constant $c$ for a given velocity $V_{Int}$ (Fig. S4E). Based on these calculations the model used a DNA constant of $c = 500$.

7. Low Entrainment Region

The RNAP convoy model is justified by the energetic cost of spinning the DNA, friction, and the ratio between topoisomerases and all three eukaryotic RNAPs. Theoretically, only two flanking topoisomerases might be sufficient act as swivels to release torsion generated from DNA rotation by an entire convoy of RNAP. Notably, experimental data demonstrated that depletion of both topoisomerases causes severe perturbation in rDNA transcription when RNAPI is around 2 Kb into the transcription unit (El Hage et al., 2010). This was shown using a range of methods including northern hybridization, ChIP and chromatin spreads. Decreased Top1 activity is accompanied by an increased number of R-loops, as also observed in the human rDNA (Manzo et al., 2018).

We interpret this observation as showing that RNAPI molecules are initially able to spin around the DNA, allowing changes in their relative positions without generating torsion, but become locked by torsion at around +2 Kb. To incorporate this mechanism into the model, we progressively engaged torsion within RNAP convoys over the initial 2 Kb of the rDNA. We used a linear engagement scheme, where at position 0 RNAP moves according to discrete stochastic elongation, at position 1000 DNA torque was applied in 50% and becomes fully engaged at position 2000 and later. This Low Entrainment Region was implemented as three elements: (1) Decreased DNA stiffness constant $c$. (2) Reset of $x_{Init}$ position. (3) A small decrease in the intrinsic RNAPI velocity (≤ 20%) to mimic the cost of friction. All three elements were applied progressively.

8. Role of nascent RNA in transcription elongation

Finally, we introduced our findings on the effects of sequence in the RNA:DNA hybrid within the transcription bubble and the structure of the extruded RNA into the model.

Nascent RNA interacts with template strand of DNA within transcription bubble. Stronger hybrids usually have a higher G+C content, particularly with G in the DNA sequence, and this correlates with slower RNAP translocation. We calculated the ΔG of RNA:DNA hybrids over an 8 nt rolling window ($dG^{RNA:DNA}$) along the rDNA as previously described (see; (El Hage et al., 2014; Turowski et al., 2016)).

The folding energy of nascent RNA was calculated using a 65 nt rolling window, offset by 15 nt ($dG^{Structure}$), as described in Materials and Methods. Stronger structures limit translocation backward and promote translocation forward. Hence, RNA structures adjacent to RNAPI would act on elongation rate positively. On the basis of the backtracking assay (Fig. 3 and 4) we applied this parameter only for structures with folding energy below the threshold value ($\Delta G \leq -11$ kcal·mol$^{-1}$). This excluded an artificial situation when long, but very weak structures would apparently have a sufficiently low ΔG to promote translocation.

Both values were incorporated into a model as modifiers of RNAP jump probability as follows:

$$V = V_{Random+Torque} + dG^{Structure}_{Strength} \cdot dG^{Structure} - dG^{RNA:DNA}_{Strength} \cdot dG^{RNA:DNA}$$

Values of strengths were fitted. Further details are in the model optimization section.

9. Optional elements of the model

A number of additional factors were considered during development of the model:

a) Topoisomerase activity

   In our model topoisomerases induce single-strand cuts to spin DNA when a convoy of RNAP generates sufficient rotating force. The canonical role of topoisomerases is associated with resolving DNA supercoiling and we tested this possibility. Top1 can unwind a minimum of one complete turn of DNA. Therefore, we applied Top1 activity as a probability function of resolving a complete turn when distance between adjacent RNAP particles was greater than 25 nt. As demonstrated on Fig. S4F Top1 activity has minor effect on the overall profile.

b) Premature termination

A potential explanation for the 5' bias in the RNAPI CRAC profile was premature termination. RNAPII is known to undergo transition from initiation state to elongation state that is associated with changes of phosphorylation status of C-terminal (Milligan et al., 2016). We considered that RNAPI might undergo a similar transition, with the region of the 5' bias reflecting a region in which RNAP has an elevated probability to terminate. Application of premature termination recapitulates the overall shape of the profile but greatly reduces the total number of RNAP per transcription unit (Fig. S4G and S4H). We were unable to find a probability where both criteria, (i) overall profile and (ii) number of RNAP molecules per rDNA, were satisfied. Matching the 5' bias was accompanied by a 30% lower number of RNAPI molecules per rDNA than observed using Miller spreads. Nevertheless, cannot exclude premature termination of RNAPI or at least partially playing role in establishing the 5' bias. However, from our modeling, it does not appear to be a key factor.

c) R-loops

R-loops arise when nascent RNA hybridizes with melted DNA helix and constrain progression of RNAP. To include r-loops as a parameter of transcription elongation model their length and position would have to be established. The distribution of RNA-DNA hybrids has been mapped, genome-wide by methods using anti-RNA:DNA antibody (El Hage et al., 2014; Wahba et al., 2016). The median length of r-loop prone genomic regions in yeast was reported to be 500 nt (Wahba et al., 2016), but is unlikely to be the length of actual DNA:RNA hybrids within the rDNA region. Nascent pre-rRNA is co-transcriptionally bound and processed by a multi-protein complex, so called small subunit processome (Turowski and Tollervey, 2015). Only short fragments of free, nascent pre-rRNA are expected to be available for potential RNA:DNA hybrid formation, making the availability of single stranded, nascent RNA rate-limiting. This availability will be anti-correlated to folding energy of nascent RNA extruded from the RNAP; i.e. hairpins within the 5'ETS region should also reduce formation of r-loops. In consequence, the potential for r-loop formation is indirectly implemented into the model by a RNA folding element and there is no need to introduce an additional factor.

*Numerical convergence of the model*

The stochastic model contains three numerical parameters: the time step, the total time for each independent simulation and the number of independent simulations to be averaged.

The principle constraint on the time step is that the distance moved in a single time step should be 0, 1 or -1 (since only single nucleotide jumps are permitted). As an estimate, we note from Fig. S4B' that the probability of sampling a velocity larger than 120 $nt \cdot sec^{-1}$ is very small. We hence take an initial time step estimate of 1/120 ~ 0.008. We performed test simulations at this and half the time step (0.004) and noted that there were no significant differences. All remaining simulations were performed with this time step.

We determined the total time necessary to run the model by monitoring expected values, such as the number of particles and the mean separation, requiring that these had reached equilibria. The main purpose of this was to remove bias caused by initiation of the RNAP molecules along the transcription unit. We found no significant differences when the total time was between 1500 sec and 3000 sec.

Increasing the number of independent simulations decreases the statistical noise in the final result. This can also be achieved by increasing the total time of each simulation, but due to parallelization, it is more efficient to increase the number of simulations. We performed convergence studies for a range of parameters and determined that there is no significant difference between results with 256, 512, and 1024 independent simulations. For the parameter studies below, due to the large number of parameter combinations, we used 256 simulations, whereas for the single chosen parameter set we used 1,024.

*Model optimization*

The model was optimized towards two major criteria: (1) The number of RNAPI molecules present on the transcription unit (Fig. S4I). (2) The general shape of the occupancy plot relative to that obtained with CRAC (Fig. 1D).

Given the constraints on the parameters discussed above, we tested all parameter combinations with transcription initiation (addProb) = {0.7,0.8,0.9}11, DNA stiffness constant *c* = {400,500,600}, $dG_{Strength}^{Structure}$ = {1,1.25,1.5}, $dG_{Strength}^{RNA:DNA}$ which was represented as a ratio to $dG_{Strength}^{Structure}$, with ratio = {0.32,0.48,0.64}, threshold value of folding energy (structure2consider) = {-10,-11,-12}. This gave us a total of $3^5$ = 243 sets of parameters, with each varying approximately 10-20% from the chosen value. Figure S4J demonstrates that the main features of the results (shape and position of peaks, general profile, number of particles), are robust under these variations in the parameters. We also demonstrate that the chosen parameters give a representative data set, lying approximately in the middle of the set of simulations over all parameters.

*Data sampling*

In order to mimic the experimental measurement process, we applied a smooth cutoff function to the data, essentially reducing the measurement of RNAPI in areas of low density/high velocity. The cutoff function is given by

$$rho_{Exp} = cutOff \cdot rho$$

$$cutOff = 0.5[1 + \mathrm{erf}\left(\frac{(rho - rho_0}{\sigma}\right)]$$

where $rho_0$ and $\sigma$ are parameters that determine the $cutOff$ position and range. We note that the *in silico* density profiles are normalized so that they have unit area; they are probability distributions. To maintain this, the 'experimental' densities are renormalized after the $cutOff$.

*Relative contribution of model elements*

To calculate the relative contributions of different forces to the modeled elongation, absolute values were used. RNA structures always act positively, RNA:DNA hybrids act negatively, whereas DNA Torsion can act both, positively or negatively. All three modifiers were summed for each nucleotide position and their relative contributions were calculated as a percent of that sum.

### Supplementary Figures

#### Supplementary Figure 1

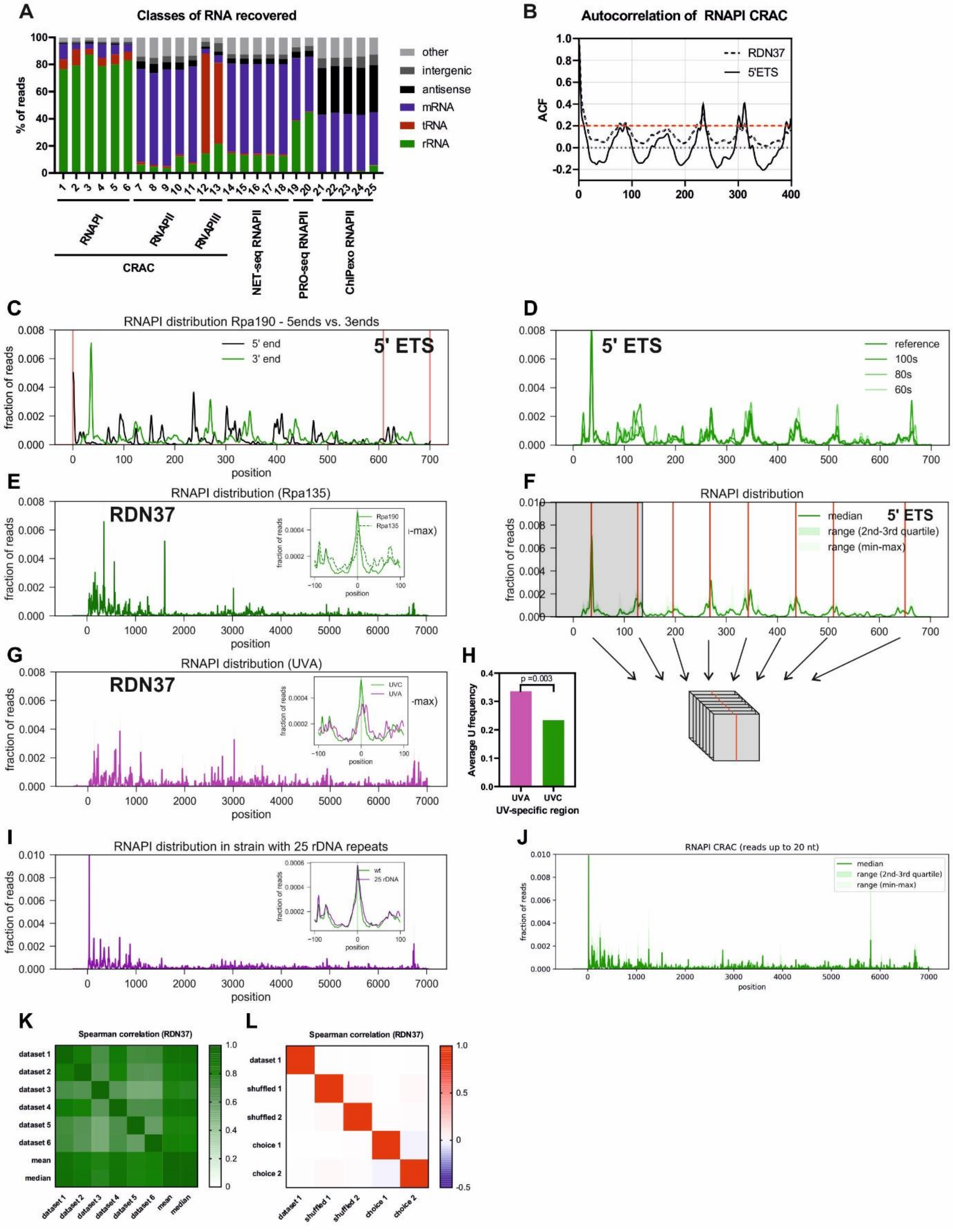

**Figure S1.** Initial analysis of RNAPI CRAC data.

(A) Transcriptome-wide binding profiles for RNAP I, II and III from CRAC (lanes 1 to 13), RNAPII NET-seq (lanes 14-18), PRO-seq (lanes 19-20) and ChIPexo (lanes 21-25).

(B) Autocorrelation function for the 5' ETS region of the pre-rRNA (solid line) and entire *RDN37* transcription unit (dashed line) indicates ~80 nt spacing between peaks in RNAPI CRAC data.

(C) RNAPI CRAC profile over the 5'ETS. The 3' ends (median, green) and the 5' ends (median, black) of mapped RNAs were plotted. There is no clear enrichment of reads at cleavage sites A0 and A1 (red lines).

(D) Rpa190-HTP CRAC profiles for UVC cross-linking time course. The 5' ETS region is presented to show data reproducibility for 60, 80 and 100 seconds of UVC cross-linking time.

(E) Rpa135-HTP CRAC profile, with the second largest RNAPI subunit tagged. Embedded: RNAPI CRAC peak metaplot for *RDN37*, comparing Rpa190 and Rpa135 at the level of single peaks.

(F) Outline of peak metaplot generation. All peaks were found using a dedicated function (see Materials and methods). Windows around each peak were superimposed to generate a metaplot centered around the peaks.

(G) Rpa190-HTP PAR-CRAC (UVA) profile. Embedded: Rpa190-HTP CRAC (UVC) peaks metaplot for *RDN37* overlaid with Rpa190-HTP PAR-CRAC (UVA). The data shows reproducibility on a level of single peaks.

(H) Average U frequency is slightly higher at the 3' end of reads recovered with PAR-CRAC. U frequency was calculated for the last two nucleotides for regions specifically enriched with UVA (purple) or UVC (green). P-value was calculated using a two-sided T-test.

(I) Rpa190-HTP CRAC profile for *RDN37* obtained from strain with 25 rDNA repeats. Embedded: Rpa190-HTP CRAC peaks metaplot for *RDN37* comparing wt and 25 rDNA repeat strains at the level of single peaks.

(J) Rpa190-HTP CRAC profile for *RDN37* obtained from reads only up to 20 nt.

(K) Spearman correlation between datasets 1-6, and median or mean of these datasets.

(L) Spearman correlation between dataset 1 and randomized datasets.

### Supplementary Figure 2

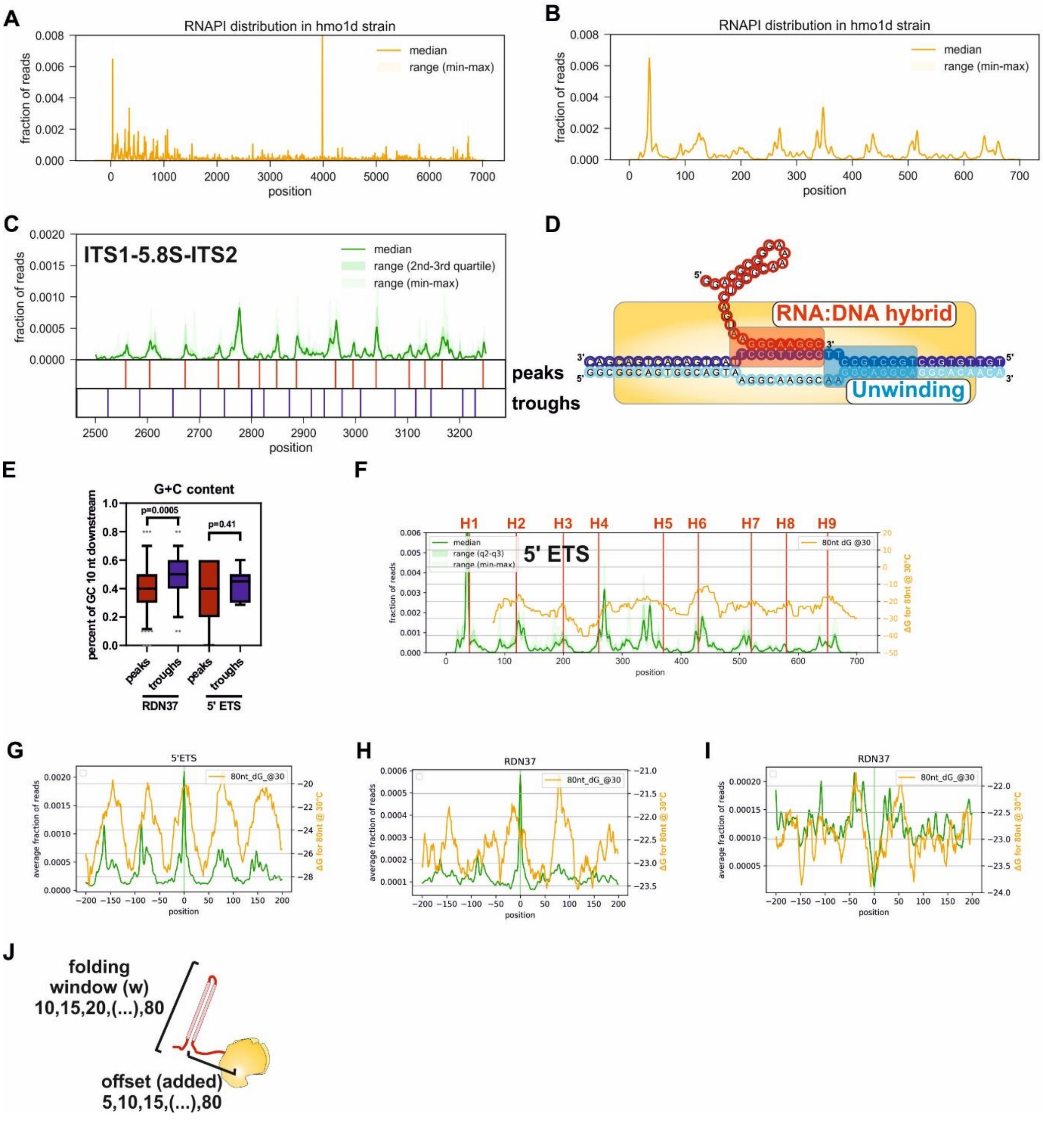


**Figure S2.** RNAPI density correlates with features in the nascent pre-rRNA.

(A) RNAPI CRAC profile for *RDN37* from the Rpa190-HTP *hmo1Δ* strain. The peak at 4000 nt is an artefact previously observed in CRAC analyses.

(B) RNAPI CRAC profile for 5' ETS from the Rpa190-HTP *hmo1Δ* strain.

(C) RNAPI CRAC profile for the ITS1, 5.8S rRNA and ITS2 regions with marked peaks and troughs used for further analysis.

(D) Schematic representation of nucleotides considered to play role in RNA:DNA hybrid formation (red box) or DNA unwinding (blue box).

(E) G+C-content in front of RNAPI has a minimal impact on DNA unwinding. Boxplot presenting distribution of G+C-content among peaks and troughs within RDN37 or the 5' ETS. G+C content is calculated for 10 nt downstream from each feature.

(F) RNAPI CRAC profile (green) for the 5' ETS plotted together with known hairpins within the 5' ETS structure (red, vertical lines with helix number above) and folding of the nascent transcript using different 80 nt folding windows at 30°C. Folding energy was calculated using hybrid-ss-min from the UNAfold package.

(G) RNAPI CRAC peaks metaplot for the 5' ETS with folding energy of the nascent transcript (80 nt folding window). *Note*: Stronger structures have lower ΔG.

(H) RNAPI CRAC peaks metaplot for *RDN37* with folding energy of nascent transcript (80 nt folding window).

(I) Same as (H) but troughs were superimposed.

(J) Schematic representation of all parameters tested for calculation of folding window (10-80 nt) and offset (5-80 nt).

### Supplementary Figure 3

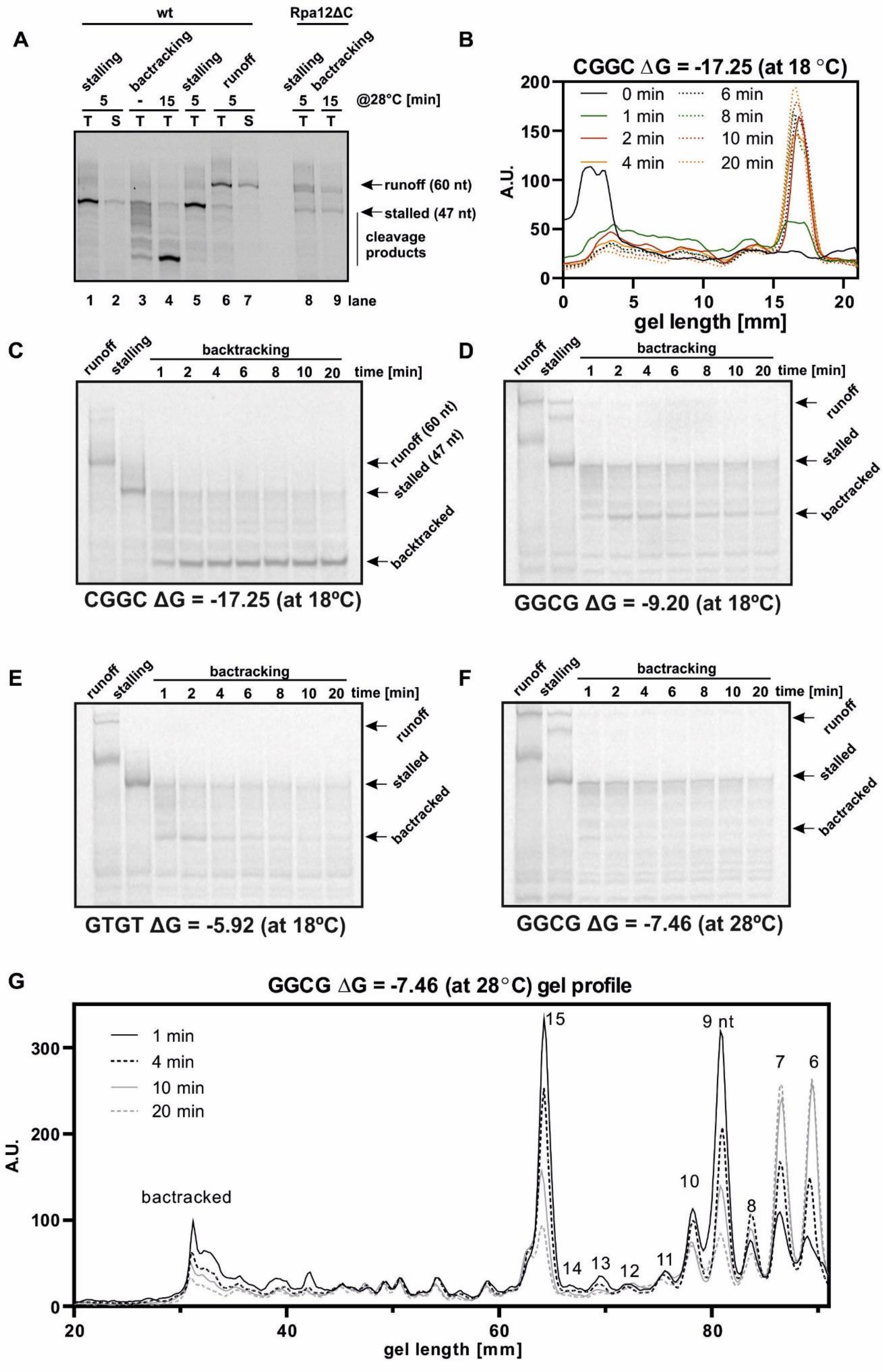

**Figure S3.** Nascent RNA limits backtracking proportionally to folding energy.

(A) RNAPI cleavage requires the Rpa12 C-terminal domain. *In vitro* assay for RNAPI backtracking, performed after initial incubation with transcription buffer (TB) lacking ATP ("stalling"). Nucleotides were washed out and samples incubated in TB without NTPs to induce RNAPI backtracking ("backtracking"). Cleavage products were detectable for RNAPI with wt Rpa12, but were not observed with Rpa12ΔC. Unexpectedly, the C-terminal truncation of Rpa12 allowed substantial readthrough of the AAA stalling sequence during extension in the absence of added ATP (stalling, lane 8). This observation will be followed up elsewhere.

(B) Quantification of all lanes from Fig. 3A presenting area between stalled and backtracked peaks.

(C) – (F) Representative gels showing backtracking assays used to study kinetics of RNAPI translocation. Assay performed as in Fig. 3C. The scaffold used, temperature and predicted folding energy are indicated below each panel.

(G) Quantification of backtracked peaks and shorter products from backtracking assay. For clarity, only four time points are displayed. The scaffold used, temperature and predicted folding energy are indicated above the plot.

### Supplementary Figure 4

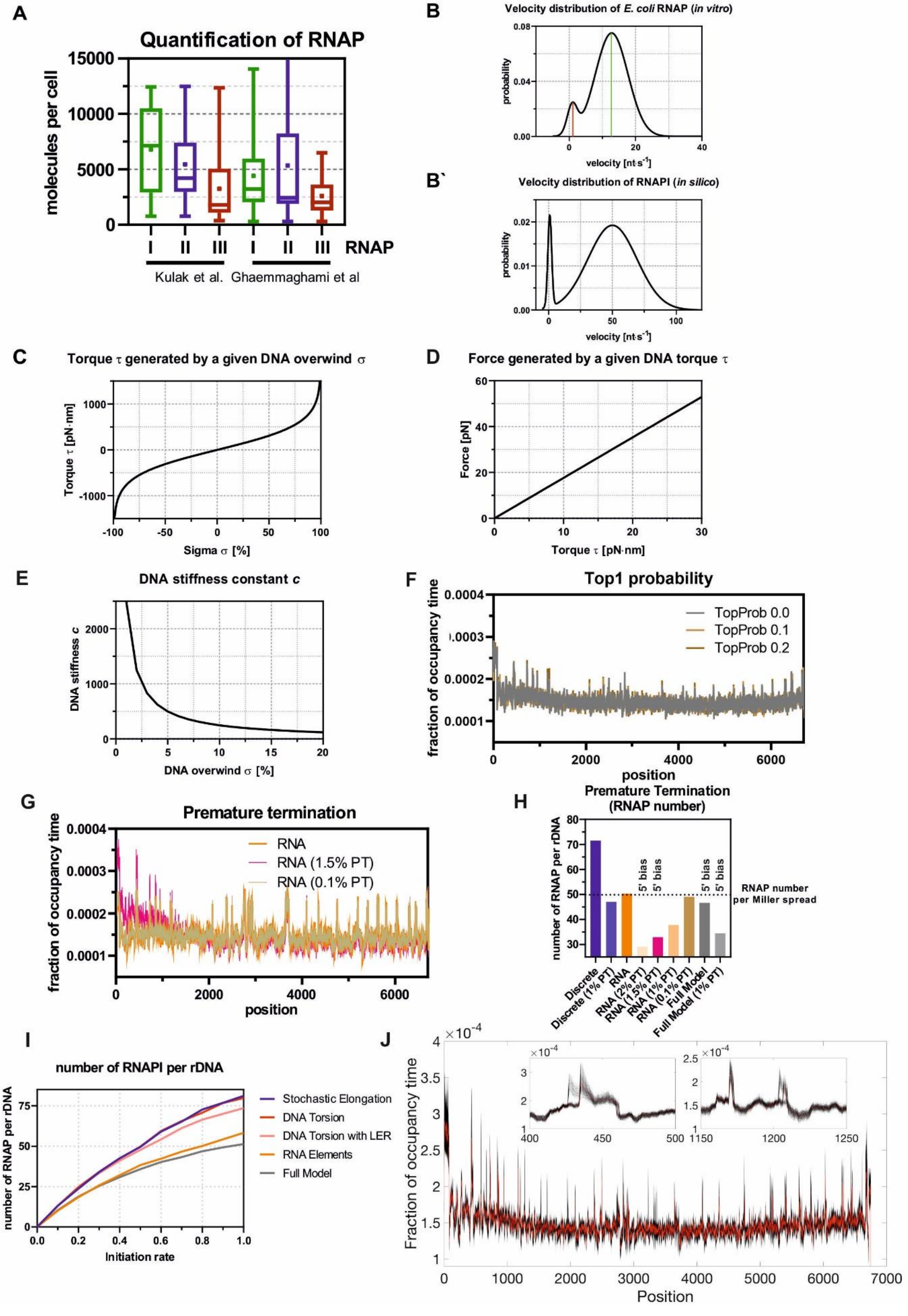

**Figure S4.** Mathematical model of RNAPI transcription.

A: Quantification of RNA polymerases in yeast cell. Boxplots for each RNA polymerase using two independent studies and all detected subunits.

B: Velocity distribution of bacterial RNAP (Adelman et al., 2002). Described by two Gaussian functions: reflecting 7.8% of the area, represents paused state and is centered at 0.9 $nt{\cdot}s^{-1}$ and a second function, which reflects active elongation and is centered at 12.8 $nt{\cdot}s^{-1}$.

B': Same as B, but the active elongation function is centered at 50 $nt{\cdot}s^{-1}$.

C: Function of DNA torque $\tau$ generated by a given DNA overwind $\sigma$.

D: Function of force $F$ generated by a given DNA torque $\tau$.

E: Function of DNA stiffness constant $c$ in relationship to DNA overwind $\sigma$ for $V_{Int}$=50 $nt{\cdot}s^{-1}$.

F: Effect of Top1 (n=64). The results of simulations using parameter TopProb = {0, 0.1, 0.2} were superimposed to show high similarity of overall profile.

G: Effect of premature termination (n=64). The results of simulations allowing for premature termination over first 2000 nt with a given probability = {0%, 0.1%, 1.5%} were superimposed to show high similarity of overall profile.

H: Number of RNAPI molecules per rDNA relative to the premature termination rate (panel Fig. S5G) Note: The 5' bias of the profile (panel I) is accompanied by ~35% drop of RNAP molecules, whereas stable number of RNAPI molecules do not recapitulate the 5' bias (panel J).

I: Number of RNAPI molecules per rDNA relative to the initiation rates are shown for different version of the model used in Figs. 4B-4G.

J: Parameter Fitting (simulations n=256). The results for each parameter set are plotted in (transparent) grey; darker areas indicate more likely results. The red line denotes our choice of parameters (simulations n=1,042). Insets show two zoomed areas. Note that the results are robust under changing parameters, and the results presented for the chosen parameter set are 'typical'.

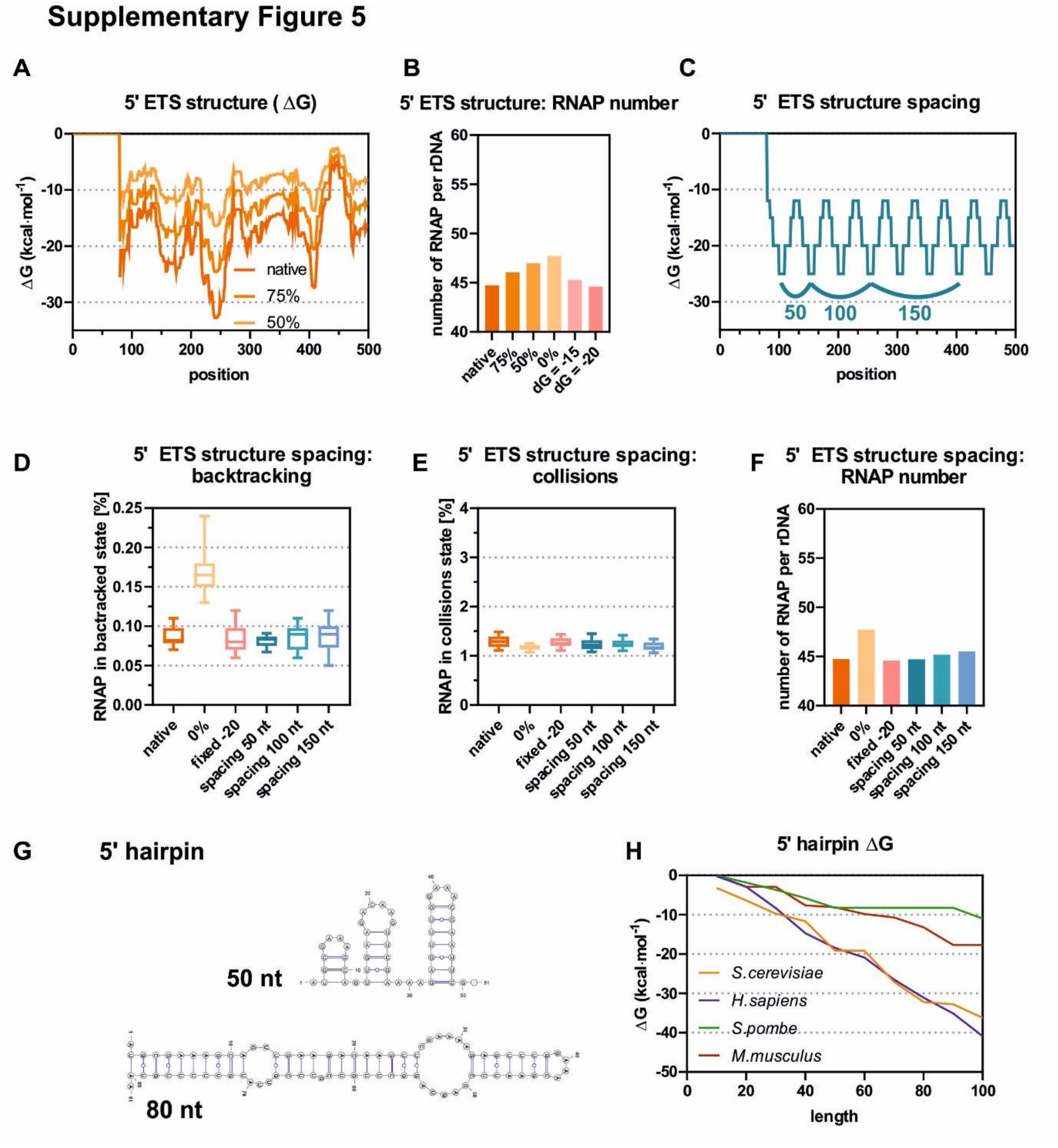


**Figure S5.** Conclusions from the model on the role of 5’ ETS structure.

A: Plot of the folding input data for the first 500 nt. Folding energy of the 5’ ETS was calculated for a rolling window of 65 nt with 15 nt offset (dark orange). Two modified input files are presented, with folding reduced to 75% or 50% of the biological values. The input with a fixed ΔG would be a straight line and is omitted for clarity.

B: The predicted number of RNAPI molecules loaded on the rDNA remains similar for altered 5’ ETS structures.
C: Overview of altered spacing of 5’ ETS hairpins. Structural data were generated as presented with peak spacing equal 50. For 100 and 150 nt spacing, peaks were connected using longer spacers of fixed energy $\Delta G = -12\ kcal\cdot mol^{-1}$.
D-F: Modification of spacing between hairpins within the 5’ETS. Measured features of the model remain unaffected: frequency of backtracking (D), frequency of collisions (E) and RNAP number per rDNA unit (F).
G: Secondary structures of the 5’ proximal hairpin in the 5’ ETS (lower panel) and the 50 nt long transcription intermediate (upper panel).
H: Folding energy of the full-length 5’ proximal hairpins and transcription intermediates. The dG was calculated for first 100 nt of the 5’ ETS and for shorter intermediates at 10 nt intervals, using sequences from *S. cerevisiae, H. sapiens, S. pombe* and *M. musculus*.

### Supplementary Figure 6

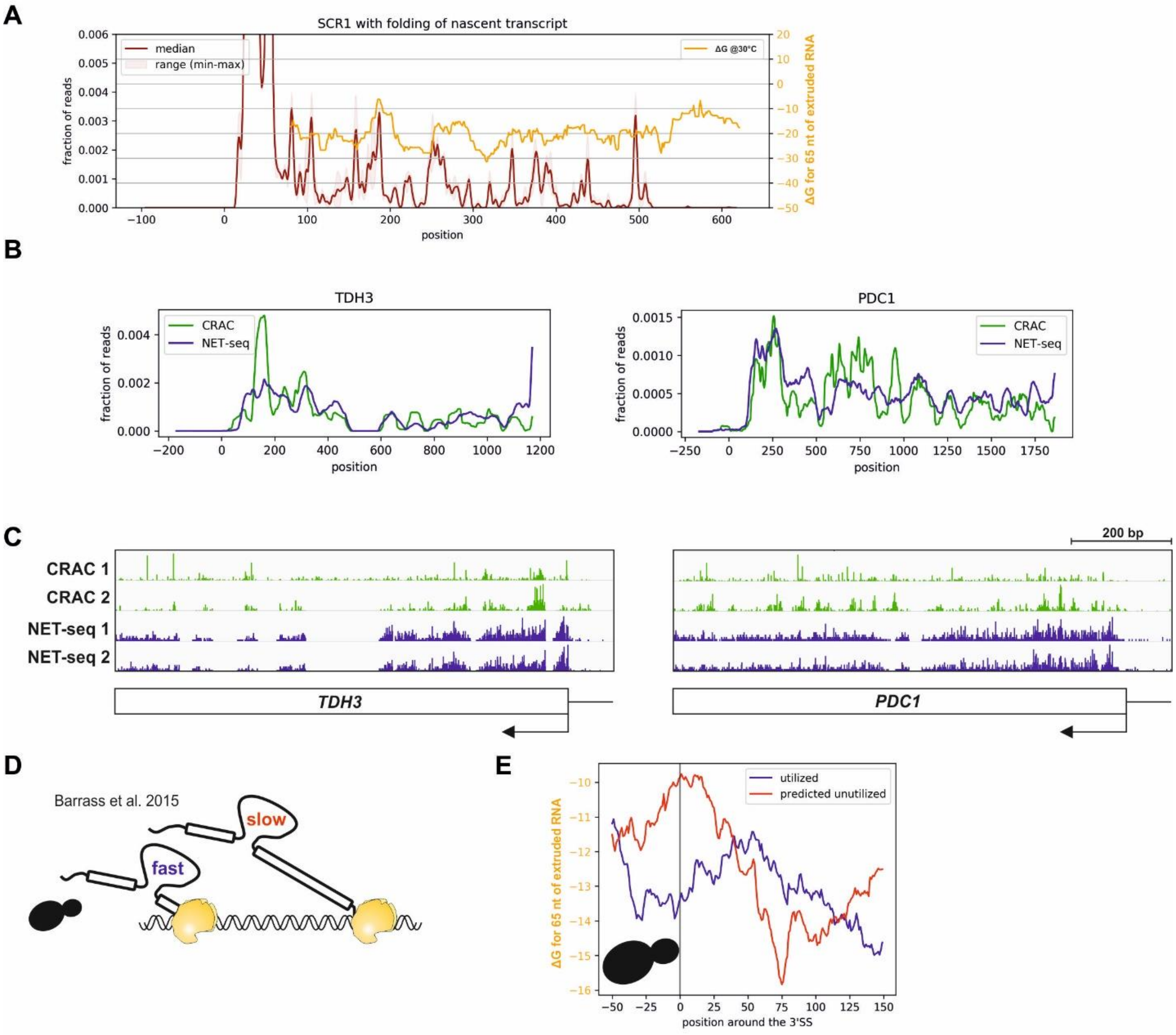


**Figure S6.** Folding of the nascent transcript plays a role in determining elongation rate of all eukaryotic RNA polymerases.

A: RNAPIII profile for *SCR1* overlaid with nascent RNA folding. The same folding window as for RNAPI analysis was used (65 nt rolling window with 15 nt offset).

B: RNAPII density across *TDH3* (left panel) and *PDC1* genes (right panel). Medians of NET-seq (blue) and CRAC (green) data were plotted to show similarities between the profiles.

C: Genome Browser tracks for RNAPII CRAC (green) and NET-seq (blue) for two highly expressed genes *TDH3* and *PDC1*. Two replicates are shown.

D: Cartoon indicating differences between fast and slow-spliced pre-mRNAs. Slow splicing could be also considered as post transcriptional.

E: Folding energy of nascent RNA around utilized 3’ SS, versus predicted but skipped 3’ SS. The same data were used to generate Fig. 6J. Note that folding of the nascent RNA does not affect accessibility of the 3’ SS.
